# Chloroplasts lacking class I glutaredoxins are functional but show a delayed recovery of protein cysteinyl redox state after oxidative challenge

**DOI:** 10.1101/2023.10.31.564817

**Authors:** Finja Bohle, Jacopo Rossi, Sadia S. Tamanna, Hannah Jansohn, Marlene Schlosser, Frank Reinhardt, Alexa Brox, Stephanie Bethmann, Stanislav Kopriva, Oliver Trentmann, Peter Jahns, Marcel Deponte, Markus Schwarzländer, Paolo Trost, Mirko Zaffagnini, Andreas J. Meyer, Stefanie J. Müller-Schüssele

## Abstract

Redox status of protein cysteinyl residues is mediated via glutathione (GSH)/glutaredoxin (GRX) and thioredoxin (TRX)-dependent redox cascades. An oxidative challenge can induce post-translational protein modifications on thiols, such as protein *S*-glutathionylation. Class I GRX are small thiol-disulfide oxidoreductases that reversibly catalyse *S*-glutathionylation and protein disulfide formation. TRX and GSH/GRX redox systems can provide partial backup for each other in several subcellular compartments, but not in the plastid stroma where TRX/light-dependent redox regulation of primary metabolism takes place. While the stromal TRX system has been studied at detail, the role of class I GRX on plastid redox processes *in vivo* is still unknown. We generate knockout lines of *GRXC5* as the only chloroplast class I GRX of the moss *Physcomitrium patens*.

While we find that class I PpGRXC5 has high activities in glutathione-dependent oxidoreductase assays using hydroxyethyl disulfide or redox-sensitive GFP2 (roGFP2) as substrates *in vitro*, Δ*grxc5* plants show no detectable growth defect or stress sensitivity, in contrast to mutants with a less negative stromal *E*_GSH_ (Δ*gr1*). Using stroma-targeted roGFP2, we show increased protein Cys oxidation and decreased reduction rates after oxidative challenge in Δ*grxc5* plants *in vivo*, indicating kinetic uncoupling of the protein Cys redox state from glutathione redox potential. Protein Cys disulfide and *S*-glutathionylation formation rates after H_2_O_2_ treatment remained unchanged. Lack of class I GRX function in the stroma did not result in impaired carbon fixation.

Our observations suggest specific roles for class I GRX in the efficient redox equilibration between *E*_GSH_ and protein Cys in the plastid stroma as well as negligible cross-talk with metabolic regulation via the TRX system. We propose a model for stromal class I GRX function as efficient kinetic couplers of protein Cys redox state to the dynamic stromal *E*_GSH_ and highlight the importance of identifying *in vivo* target proteins of GRXC5.

**One sentence summary:** Removal of class I GRX activity in the chloroplast stroma of *P. patens* kinetically uncouples GRX-dependent cysteine redox changes from the local glutathione redox potential, without an effect on NPQ or photosynthetic carbon reactions.

## Introduction

Oxygenic photosynthesis has shaped our planet by increasing oxygen levels in the atmosphere, and by enabling solar-driven carbon fixation. In phototrophic eukaryotic life forms, former free-living cyanobacteria now serve as chloroplasts in light-harvesting and production of reducing equivalents to power reductive processes in cell metabolism (Schreiber et al., 2022). Land plant chloroplasts face multiple oxidative challenges as environmental conditions such as light intensity, water availability and temperature can rapidly fluctuate and cause imbalances between light reactions and carbon fixation. Plants have adapted to a frequently changing environment by evolving mechanisms to regulate photochemistry and carbon fixation in a matter of minutes, as well as mechanisms to acclimate to a changed steady state in a matter of hours or days (Roach and Krieger-Liszkay, 2014; Schöttler and Toth, 2014; Alboresi et al., 2019; Morales and Kaiser, 2020). Regulation of the enzymatic reactions of the Calvin-Benson-Bassham cycle (CBB cycle) to match the activity of the light reactions guarantees efficient re-oxidation of the electron acceptor of light reactions, NADP^+^, and avoids futile cycling under dark conditions when the oxidative pentose phosphate pathway is active (Buchanan et al., 2012). Thus, many stromal enzymes evolved Cys-based redox-regulation (Buchanan and Balmer, 2005; Balsera et al., 2014). This regulation of protein activity or oligomerization (Marri et al., 2014) can be mediated by post-translational changes in cysteinyl thiol redox states, including the formation/reduction of regulatory disulfide bonds on target proteins (Michelet et al., 2013; Gütle et al., 2016; Gurrieri et al., 2023). The thioredoxin (TRX) system derives electrons from the photosynthetic electron transport (PET) or NADPH for the reduction of specific regulatory disulfides on metabolic enzymes (Geigenberger et al., 2017). The redox state of these thiol-switches depends on the redox state of TRXs, which in turn depend on PET/NADPH-dependent reduction rates (Zimmer et al., 2021; Teh et al., 2023) and oxidation rates (Yoshida et al., 2019). TRX oxidation rates can be linked via 2 Cys-peroxiredoxins (PRX) to the detoxification of hydrogen peroxide (H_2_O_2_) (Pérez-Ruiz et al., 2017; Ojeda et al., 2018; Vaseghi et al., 2018) which functions as terminal electron acceptor.

When the stromal NADP pool becomes increasingly reduced, generation of reactive oxygen species (ROS) increases (photoinhibitory conditions) as photosystem I (PSI) becomes acceptor-limited (Roach and Krieger-Liszkay, 2014). Via superoxide dismutases, superoxide is rapidly (without input of additional electrons) converted to H_2_O_2_ that can react with cysteine residues causing thiol oxidation, either directly (highly reactive Cys only) or via the Cys-redox relays (Deponte, 2017; Alvarez and Salinas, 2022). To balance ROS formation and repair ROS-induced oxidative damage, chloroplasts have evolved multiple detoxification and repair systems that either draw electrons from the TRX system or the glutathione system.

Glutathione (GSH) is a cysteine-containing tri-peptide present at millimolar concentrations in the cytosol and chloroplast stroma serving multiple roles in cellular metabolism and defense (Bangash et al., 2019; Cassier-Chauvat et al., 2023). First, glutathione is an important electron donor for H_2_O_2_ detoxification or repair of ROS-induced damages (reviewed in Noctor et al., 2012 and Müller-Schüssele et al. (2021a)). Second, it can form mixed disulfides with cysteinyl residues in proteins (protein *S*-glutathionylation), either as a consequence of cysteine oxidation by H_2_O_2_ (*S*-sulfenylation), or enzymatically catalysed by class I glutaredoxins (GRXs) (**Fig. 1**) (Deponte, 2017). Class I GRXs are small oxidoreductases belonging to the TRX superfamily, that can form or release protein *S*-glutathionylation and disulfides (Fernandes and Holmgren, 2004; Lillig et al., 2008; Couturier et al., 2013; Trnka et al., 2020). *S*-glutathionylation can affect protein activity and/or oligomerization as well as act as a protection against protein Cys over-oxidation, depending on the modified protein and site (Michelet et al., 2005; Zaffagnini et al., 2007; Bedhomme et al., 2009; Zaffagnini et al., 2013). GRXs and TRXs can at least partially functionally complement each other, lack of both activities in the cytosol leads to lethality in *S. cerevisiae* (Draculic et al., 2000). Thus, the redox state of individual cysteine residues depends on an intricate network of redox reactions with partially overlapping but also specific roles for TRX, NTRC- and GSH/GRX-dependent reactions.

**Figure 1:**
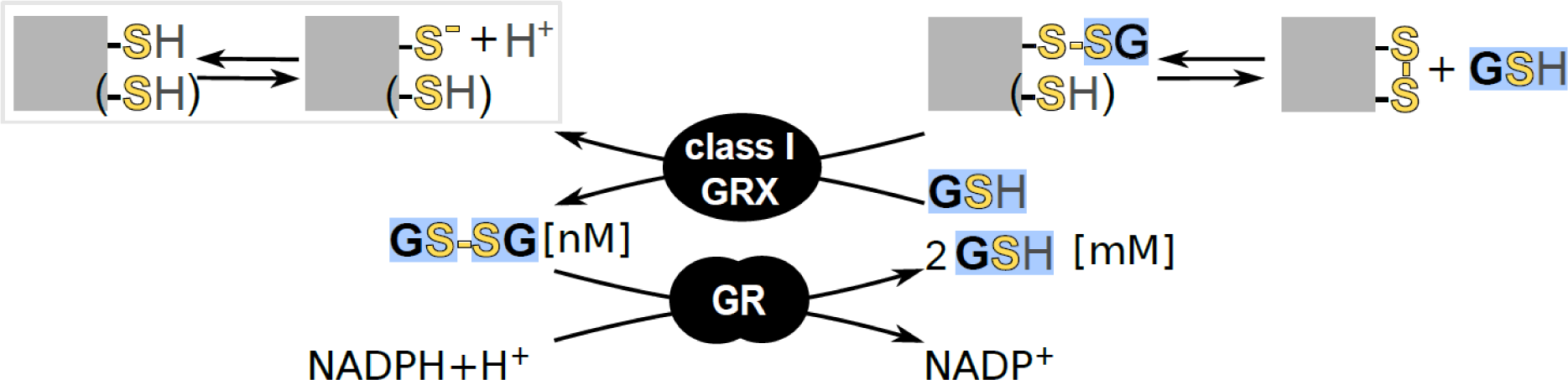
Scheme of class I GRX and glutathione reductase function. Class I glutaredoxins (GRX) can reversibly modify cysteinyl residues (thiol group SH, thiolate S^-^) in proteins by forming a mixed disulfide with the tripeptide glutathione (GSH). If a second cysteine is present in a suitable distance, this *S*-glutathionylation can be released by intramolecular disulfide formation, as observed for the genetically encoded redox sensor roGFP2. Glutathione reductase (GR) regenerates glutathione disulfide (GSSG) to 2 glutathione (GSH) at the expense of NADPH, keeping the glutathione redox potential *E*_GSH_ reducing for most cysteines. Grey squares represent GRX substrate proteins, black rounded shapes enzymes.

In the plastid stroma, the presence of a glutathione reductase (GR) leads to a highly reducing steady-state glutathione redox potential (*E*_GSH_) with only nanomolar amounts of oxidized glutathione, glutathione disulfide (GSSG) (**Fig. 1**) (Yu et al., 2013; Schwarzländer et al., 2016; Marty et al., 2019; Müller-Schüssele et al., 2020). Stromal steady-state *E*_GSH_ monitored by the genetically encoded biosensor Grx1-roGFP2 (Meyer et al., 2007; Gutscher et al., 2008; Schwarzländer et al., 2008) revealed light-dependent redox dynamics (Müller-Schüssele et al., 2020; Haber et al., 2021).

In contrast to mitochondria and cytosol, the plastid TRX system does not constitute an effective functional backup system for the stromal GSH/GRX system, as a lack in stromal GR causes embryo-lethality in *A. thaliana* (Marty et al., 2009; Marty et al., 2019). In the model moss *Physcomitrium patens*, plants lacking mitochondria/plastid-targeted glutathione reductase (PpGR1) had a shifted (less negative) stromal *E*_GSH_ and were viable, albeit dwarfed and light-sensitive (Müller-Schüssele et al., 2020). Plant class I GRX clades that contain isoforms targeted to different subcellular compartments are evolutionary conserved from bryophytes to flowering plants (Müller-Schüssele et al., 2021a). In the model flowering plant *A. thaliana*, two plastid-targeted class I GRXs exist that differ by the number of cysteines in the active site. While AtGRXS12 contains a single cysteine (WCSYS active site), AtGRXC5 contains two cysteines (YCPYC active site) (Couturier et al., 2011; Müller-Schüssele et al., 2021a). According to previous phylogenetic analysis, GRXC5 represents the ancestral type of plastid-targeted class I GRX with a single isoform of GRXC5 in the model moss *P. patens* (Müller-Schüssele et al., 2021a).

The chloroplast stroma has been described as a ‘redox battle ground’ (Meyer et al., 2021) of which we are still lacking a functional map. In particular, the roles of glutathione, class I GRX and protein *S*-glutathionylation are largely uncharted. It is an open question how class I GRX function and *S-*glutathionylation dynamically interact with the known thiol-switching cascades in the crucial light/dark regulation of chloroplast metabolism.

Here, we set out to understand the role of class I GRX in the stromal redox network of plants. To this end we generated plant lines lacking class I GRX activity in the stroma, by exploiting the fact that only a single plastid-targeted class I GRX present in *Physcomitrium patens.* We combine biochemical characterization of PpGRXC5 *in vitro* with *in vivo* biosensing using stroma-targeted roGFP2 to dynamically monitor protein Cys redox changes after oxidative challenge.

## Material and Methods

### *In vitro* analyses of PpGRXC5

For recombinant protein expression of PpGRXC5, *P. patens* cDNA was used to amplify *GRXC5* without targeting peptide (starting at the Ala 120 codon) using the primer combination PpGRXC5_A120_F GGGGACAAGTTTGTACAAAAAAGCAGGCTTAGCAGCAGGTT CGGGG and PpGrxC5_cds_R GGGGACCACTTTGTACAAGAAAGCTGGGTGTCAACTCCTGTTTGCACCAG, adding *attB1* and *attB2* sites. The PCR-product was cloned via Gateway™ into pDONR207, verified by sequencing, and subsequently inserted into the pETG-10A vector. Recombinant proteins (roGFP2 (Meyer et al., 2007), PpGRXC5, AtGRXC1 (AT5G63030, (Rouhier et al., 2007)) were purified from transformed *E. coli* strain *Rosetta2* as described in Trnka et al., (2020) and Ugalde et al. (2021).

#### Hydroxyethyl disulfide (HED) assay

Deglutathionylation activity of PpGRXC5 was tested according to Zaffagnini et al. (2008). Prior to the assay, the concentration of GRXC5 was determined via Bradford assay (Bradford, 1976). All chemicals were dissolved in 100 mM Tris-HCl pH 7.9. A 20 mM NADPH stock was prepared, and concentration was verified via absorption measurements using the NADPH extinction coefficient of 6.23 mM^−1^ cm^−1^. To test for the GRX concentration in which the GRX shows a linear activity, HED assays were performed in 100 mM Tris-HCl, 1 mM EDTA pH 7.9 using 1 mM GSH and 0.7 mM HED, varying the concentration of the GRX from 10-50 nM. The HED assay, with GSH as variable substrate, was performed by preparing a 1 ml cuvette containing 0.5-4 mM GSH and 0.7 mM bis(2-hydroxyethyl)disulfide (HED). With HED as variable substrate, GSH concentration was kept constant at 1 mM, while HED concentrations varied from 0.3 mM to 1.5 mM. To the HED and GSH mixture 200 µM NADPH and 100 mM Tris-HCl 1 mM EDTA pH 7.9 were added. After exactly 3 min of incubation, GR (final concentration of 6 µg ml^-1^, *Saccharomyces cerevisiae*, Sigma-Aldrich CAS 9001-48-3, 100-300 units/mg protein) and GRXC5 (final concentration 30 nM) were added to the cuvette adding up to a final volume of 1 ml. For each concentration of varying GSH or HED, a background activity was determined by replacing GRX with buffer (Tris-HCl, pH 7.9). The absorbance decrease at 340 nm was followed for 1 min using a NanoDrop^TM^ 2000c spectrophotometer (Thermo Fisher Scientific).

#### roGFP2-based in vitro assays

Oxidation and reduction assays using roGFP2 (Meyer et al., 2007; Gutscher et al., 2008; Bohle et al., 2023) were performed in a 96-well plate in a fluorescence plate reader (CLARIOstar® Plus, BMG Labtech). For oxidized and reduced roGFP2 controls, roGFP2 was treated for 30 min with 10 mM DTT or 10 mM H_2_O_2_ before the assay start. To assess the reduction capacities of PpGRXC5, 1 µM of untreated (oxidized) roGFP2 was pipetted into a well containing 1 µM GRX (PpGRXC5, AtGRXC1), 100 µM NADPH and 1 unit *S. saccharomyces* GR in 100 μl of 100 mM potassium phosphate buffer pH 7.4. After measuring for 10 cycles, a final concentration of 2 mM GSH was added automatically by the injection needles of the plate reader into the respective wells. Fluorescence was followed until roGFP2 ratio stabilized. For assessing oxidation capacities of PpGRXC5, 10 µM roGFP2 was pre-reduced with 10 mM DTT for 30 min and subsequently desalted via Zeba™Spin Desalting Columns (ThermoFisher) following the manufacturer’s instructions. 1 µM of pre-reduced roGFP2 was mixed with 1 µM of PpGRXC5 or AtGRXC1 in potassium phosphate buffer pH 7.4. 2 mM GSSG was added via the injection needles of the plate reader after 5 min of initial measurements and the measurement continued until roGFP2 ratio stabilized. Fluorescence intensities were collected by excitation at 390-10 nm or 480-10 nm and emission at 530-10 nm.

### Plant materials and growth conditions

*Physcomitrium patens* (Hedw.) Bruch & Schimp ecotype ‘Gransden 2004’ (International Moss Stock Centre (IMSC, http://www.moss-stock-center.org), accession number 40001) was grown axenically and regularly sub-cultured in agitated liquid medium (KNOP medium: 250 mg l^−1^ KH_2_PO_4_, 250 mg l^−1^ KCl, 250 mg l^−1^ MgSO_4_ × 7H_2_O,1 g l^−1^ Ca(NO_3_)_2_ × 4H_2_O and 12.5 mg l^−1^ FeSO_4_ × 7H_2_O, pH 5.8) (Reski and Abel, 1985) supplemented with micro-elements (ME), (H_3_BO_3_, MnSO_4_, ZnSO_4_, KI, Na_2_MoO_4_ × 2H_2_O, CuSO_4_, Co(NO_3_)_2_). For phenotypic analyses, *P. patens* gametophores were grown on KNOP ME agar plates (12 g l^−1^ purified agar; Oxoid, Thermo Scientific, Waltham, MA, USA). Light intensity in growth cabinets was set to 70-100 µmol photons m^−2^ s^−1^ and 16:8 h light/dark cycle at 22 °C, if not indicated otherwise.

### Generation of transgenic lines

#### Δgrxc5

The complete *GRXC5* (Pp3c3_7440V3.3 (V1.6 Pp1s321_10V6.1)) coding sequence (cds) was exchanged via homologous recombination with a *nptII* resistance cassette under the control of the *NOS* promoter and terminator. The knock-out construct was assembled by triple-template PCR (Tian et al., 2004), using the following primer combinations: upstream (5’) homologous region (HR) PpGRXC5ko_5PHR_P1 ATCACAGGAAGCTATGGAAGGCA and PpGRXC5ko_5PHR_P2 TTGACAGGATCCGATAATCCCCACTTAGCACCAGG, resistance cassette GRXC5ko_npt_F: TGCTAAGTGGGGATTATCGGATCCTGTCAAACACTG and GRXC5ko_npt_R: CGTATGTGATGGCATGACAGGAGGCCCGATCTAGTA, downstream (3’) HR PpGrxC5ko_3PHR_P3 ATCGGGCCTCCTGTCATGCCATCACATACGGAACT and PpGrxC5ko_3PHR_P4 ATCTTCAGCTCCTCAGTTCCTCG (**Table S1**). *Eco*RV restriction sites were introduced by ligating the triple-template PCR product into the pJET1.2 (Thermo Scientific, Waltham, MA, USA) vector. The resulting vector was amplified in and purified from *E. coli DH5alpha* strain, digested with *Eco*RV and used for polyethylene glycol-mediated protoplast transformation as previously described (Hohe et al., 2004). Regenerating plants surviving geneticin G418 (12.5 mg l^−1^) selection for four weeks were further screened via PCR for homologous 5’ and 3’ integration of the knock-out construct at the target locus using the primer combinations GrxC5_5P_F AAGTAGGGAAAAGAGAGCACG and H3b_R CCAAACGTAAAACGGCTTGT as well as NOST_F GCGCGGTGTCATCTATGTTA and GrxC5_3P_R TGTCGTGTGTTCGGACTTCT (**Table S1**). Absence of transcript on cDNA-level was confirmed using the primer combination PpGrxC5_RT_F TTAATCGGCAGGTGTGTGGA and PpGrxC5_RT_R AAAAGCTTCTTCACGCGCAT (**Fig. S1**). Δ*grxc5* lines are available from the IMSC under the accession numbers: Δ*grxc5* #54 IMSC-Nr. 40954, Δ*grxc5* #249 IMSC-Nr. 40955). Δ*gr1* lines in Δ*grxc5* #54 genetic background were generated and identified as described in Müller-Schüssele et al. (2020), see also (**Fig. S1, Table S1**).

#### Plastid-targeted roGFP2

The construct of plastid-targeted roGFP2 was generated by overlap PCR. Two DNA templates were generated in separate PCR reactions adding *attB1* and *attB2* sites to the 5’ and 3’ ends, respectively: the plastid transketolase targeting peptide (TKTP) sequence of *N. tabacum* (Schwarzländer et al., 2008; Speiser et al., 2018) was amplified using TKTP_F (GGGGACAAGTTTGTACAAAAAAGCAGGCTATGGCGTCTTCTTCTTCTCT) and TKTP_roGFP2_R (CCTCGCCCTTGCTCACCAGCGCAGTCTCAGTT), creating an overlap to the *roGFP2* coding sequence. Similarly, *roGFP2* was amplified with TKTP-roGFP2_F (ACTGAGACTGCGCTGGTGAGCAAGGGCGAGGAG) and roGFP2-attB2_R (GGGGACCACTTTGTACAAGAAAGCTGGGTCTTACTTGTACAGCTCGTCCATG), creating an overlap with the *TKTP* sequence (**Table S1**). The overlap PCR product was cloned by Gateway BP reaction in the pDONR207 entry vector. A clone with correct sequence was recombined via LR reaction into an expression vector (*PTA2_Act5_GW*) containing a Gateway *attR1/attR2* cassette between the *PpActin5* promoter and nopaline synthase (*NOS*) terminator, as well as homologous regions for integration at the *P. patens PTA2* locus (Kubo et al., 2013; Mueller and Reski, 2015). For protoplast transformation, the expression vector was digested with *BspQ1*, cutting at the ends of *PTA2* homologous regions, and co-transformed with a second uncut plasmid containing the *hpt* resistance cassette (pJET1.2 hpt; cds of hygromycin phosphotransferase under the control of *NOS* promoter and terminator). After four weeks of selection on hygromycin (12.5 µg ml^−1^) (Mueller et al., 2014), two transgenic lines expressing plastid-targeted roGFP2 in Δ*grxc5#54* background (lines #17 (IMSC-Nr. 40957) and #21 (IMSC-Nr. 40958)) and one transgenic line in WT background (#20, IMSC-Nr. 40959) were used for further analyses and are available from the IMSC.

### Confocal laser scanning microscopy

Microscopy was carried out as described in Müller-Schüssele et al. (2020) using a LSM780 (attached to an Axio Observer.Z1) (Carl Zeiss, Oberkochen, Germany) with 25x (Plan-Apochromat 25×/0.8 Imm Korr NA0.8) or 40x (C-Apochromat 40×/1.2W Korr NA1.2) objective. roGFP2 redox state was monitored by sequential excitation at 405 nm and 488 nm, detecting emission from 508 to 535 nm. Autofluorescence was recorded using excitation at 405 nm and emission detected from 430 to 470 nm. Chlorophyll autofluorescence was monitored after 488 nm excitation at an emission of 680 to 735 nm. Image intensities and 405/488 nm ratios were calculated per pixel using a custom MATLAB (MathWorks, Natick, MA, USA)-based software (Fricker, 2016). For experiments with dark/light/dark transitions, protonema and gametophores of *TKTP-roGFP2*-expressing transgenic *P. patens* lines WT #20, Δ*grxc5* #17 and Δ*grxc5* #21 cultured in liquid medium were dark-adapted for at least 45 min. Subsequently, roGFP2 fluorescence was first imaged for 1 min in the dark (without pre-screening), then during illumination from an external light source with c. 100 µmol photons m^−2^s^−1^ from a 90° angle for 5 min (every 30 s), followed by a second period of continued imaging in the dark (Müller-Schüssele et al., 2020).

### *In vivo* plate-reader based fluorometry

Ratiometric time-course measurements for roGFP2 were conducted using a CLARIOstar® Plus plate reader (BMG Labtech). During *in vivo* time series, roGFP2 signal was detected using a sequential filter-based excitation of 400-10 nm and 482-16 nm, while emission was detected at 530-40 nm. The degree of oxidation of roGFP2 (OxD) was calculated as described in Aller et al. (2013). Protonema culture of *P. patens* expressing *TKTP-roGFP2* was dispersed and transferred to fresh KNOP-ME pH 5.8 media one week prior to measurements. 200 µl of protonema culture were pipetted with a wide cut pipette tip into wells of a 96-well plate. Cultivation media was taken up after the moss settled to the bottom of the plate and substituted by 200 µl of imaging buffer (10 mM MES, 5 mM KCl, 10 mM CaCl_2_, 10 mM MgCl_2_ pH 5.8) (Wagner et al., 2019). To conduct an *in vivo* sensor calibration, the 200 µl imaging buffer were removed with a 100 µl tip and substituted with the same volume of imaging buffer containing either 10 mM H_2_O_2_ or 5 mM DPS (2,2’-dipyridyl disulfide) for complete oxidation, and 10 mM dithiothreitol (DTT) for complete reduction.

For H_2_O_2_ recovery experiments, H_2_O_2_ in concentrations ranging from 1-10 mM were added manually to the wells. After 30 min in the respective H_2_O_2_ in concentration, buffer was exchanged to imaging buffer to follow roGFP2 re-reduction. For H_2_O_2_ oxidation rate experiments the plate reader was used in “well mode” with a cycle time of 1.55 s. 200 µl of one-week-old protonema culture expressing *TKTP-roGFP2* was transferred into a 96-well plate and pre-reduced using 10 mM DTT in imaging buffer. DTT was removed and substituted with 160 µl imaging buffer. After 60 s H_2_O_2_ was automatically injected by the plate reader to a final concentration of 2 mM.

Excitation scans were performed on *P. patens* protonema tissue expressing *TKTP-roGFP2*: 100 µl of one-week-old liquid culture was transferred into a 96-well plate and either treated for 30 min with 10 mM DTT, 5 mM DPS or imaging buffer. Wells were excited at 390-490 nm using a monochromator while emission was collected at 535-16 nm. Intensities of excitation spectra were normalized to the intensity of the isosbestic point of roGFP2 at c. 425 nm.

### CO_2_ exchange measurements

CO_2_ exchange measurements were performed with the GFS-3000 (Heinz Walz GmbH, Effeltrich, Germany). For each measurement, WT, Δ*grxc5* #54, and Δ*grxc5* #249 protonema cultures were cultivated in parallel to a similar density in liquid medium. Three days after sub-culturing, 5 ml of each culture were applied onto a nylon membrane filter with a diameter of 35 mm, placed within a miniature petri dish (Ø 42 mm). Any excess liquid was removed using a 1000 µl pipette tip. CO_2_ uptake was recorded for 7.5 min in light (75 µmol photons m^−2^ s^−1^), followed by 7.5-minute in darkness. These measurements were performed consistently at a humidity level of 98% and a temperature of 22 °C, with this cycle repeated three times per biological replicate to ensure reliability. To establish a baseline measurement, a nylon membrane filter wetted with KNOP-ME medium was used as a blank (zero point; ZP) before each measurement. The recorded ZP value was subtracted from each measurement point (MP). In **Fig. 6**, data from the last 2.5 min of the first dark cycle to the third dark cycle (dark-light-dark transition) from five biological replicates from different weeks were plotted.

### SDS-PAGE and Western blotting

Protein extraction based on chloroform-methanol precipitation was performed according to (Wessel and Flügge, 1984; Mueller et al., 2014) on frozen and pulverized plant material using the TissueLyser II (Qiagen). 100 µl lysis buffer (7.5 M urea, 2.5 M thiourea, 12.5% (v/v) glycerol, 62.5 mM Tris-HCl pH 8, 0.1% (v/v) Sigma plant protease inhibitor cocktail (P9599)) was added per 10 mg pulverized plant material. Additionally, protein thiol groups were blocked by a final concentration of freshly balanced 20 mM *N*-ethylmaleimide (NEM, Sigma 128-53-0) directly added to the lysis buffer. Protein pellets were dissolved in 50-100 µl protein resuspension buffer (50 mM Tris-HCl, 8 M urea, pH 7.5-8). SDS-PAGE samples were prepared by mixing the proteins sample with 1x non-reducing Laemmli buffer (2% (w/v) SDS, 50 mM Tris-HCl pH 6.8, 0.002% (w/v) bromophenol blue, 10% (v/v) glycerol), and size-separated using 10% Mini-PROTEAN® precast gels (Bio-Rad) (SDS-running buffer 25 mM Tris-HCl pH 8.3, 192 mM glycine, 0.1% (w/v) SDS) according to the manufacturer’s instructions, using 5 µl PageRuler™ Prestained Protein Ladder (ThermoFisher) as marker. Equal loading was confirmed by staining for >1 h in PageBlue™ protein staining solution (ThermoFisher).

Proteins were transferred to a PVDF membrane (Immobilon-P, Millipore Corporation, Billerica, MA, USA) via semi-dry Western blotting. The membrane containing the proteins was blocked in 5% (w/v) milk powder dissolved in TBS-T buffer (20 mM Tris-HCl pH 7.6, 137 mM NaCl, and 0.1% (v/v) Tween20) for 1 h at 25 °C or overnight at 4 °C before labelling with the primary antibody (α-GSH, ThermoFisher, MA1-7620, 1:1000 in 2.5% (w/v) milk powder dissolved in TBS-T). Membranes were washed three times with TBS-T for 5 min and incubated for 1 h in the secondary antibody (Goat anti-Mouse, Agrisera, AS11 1772, 1:2500 in TBS). For immunodetection the Agrisera ECL kit (Super Bright, AS16 ECL-SN) was used according to the recommendations of the supplier. Western blots were imaged using the INTAS ECL ChemoStar imaging system (Intas).

## Results

### PpGRXC5 is a class I GRX with (de)glutathionylation activity

First, we determined PpGRXC5 (de)glutathionylation activity and kinetic parameters *in vitro*, by cloning and purifying PpGRXC5 (removing the N-terminal targeting sequence, starting with Ala120). We characterized *K*_m_^app^ and *k*_cat_^app^ of PpGRXC5 using the hydroxyethyl disulfide (HED)-assay. PpGRXC5 was able to very efficiently catalyse the GSH-dependent reduction of HED (Begas et al., 2015) with catalytic efficiencies (*k*_cat_ ^app^/*K*_m_ ^app^) of 1.0 x10^5^ M^−1^s^−1^ and 2.8 x10^4^ M^−1^s^−1^ when HED and GSH were used a variable substrate, respectively (**Fig. 2A**).These catalytic efficiency constants are in a comparable order of magnitude with class I GRX7 from *S. cerevisiae*, *Plasmodium falciparum* (Begas et al., 2017) and class I GRXs from the green lineage, such as *Chlamydomonas reinhardtii* GRX1 (Zaffagnini et al., 2008), poplar PtGRXS12 (Couturier et al., 2009; Zaffagnini et al., 2012a) and AtGRXC5 (Couturier et al., 2011).

**Figure 2:**
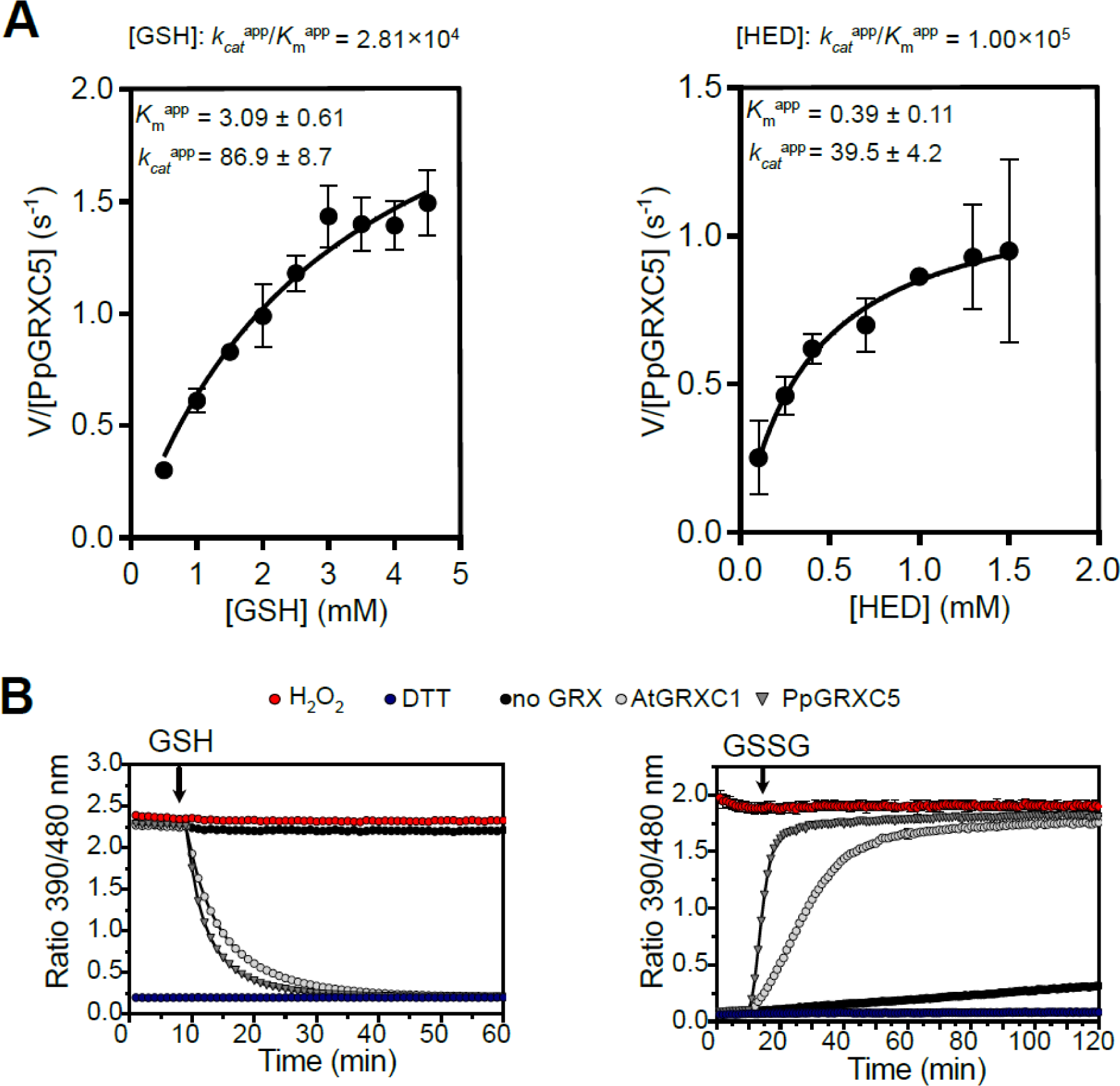
Catalytic activity of PpGRXC5 *in vitro*. **(A)** HED assays: PpGRXC5 [30 nM] was added to a cuvette containing GSH [0.5-4 mM], HED [0.3-1.5 mM], NADPH [200 μM], GR [6 μg/ml] in 100 mM Tris-HCl, 1 mM EDTA, pH 7.9. The decrease in absorbance at 340 nm was followed for 1 min (shown are means± SDs, n = 4). Varying concentration of GSH [0.5-4 mM] and a constant HED concentration of 0.7 mM was used to determine GSH-dependent kinetics (left panel). Varying concentrations of HED [0.3-1.5 mM] and a concentration of 1 mM GSH was used to determine HED-dependent kinetics (right panel). Apparent *K*_m_ (*K*_m_) is depicted in mM, apparent *k*_cat_ (*k_cat_*) in s and the rate constant (*k_cat_* / *K*_m_) in M s . Non-linear regression was fitted using Prism 9 (GraphPad). **(B)** roGFP2 reduction assay (left panel): 1 μM of PpGRXC5 or 1 μM AtGRXC1 were incubated with 1 μM of oxidized roGFP2 in 100 mM KPE, pH 7.4. Arrow indicates the time point of addition of 2 mM GSH; n = 3 ± SDs. roGFP2 oxidation assay (right panel): 1 μM of PpGRXC5 or 1 μM AtGRXC1 were incubated with 1 μM of pre-reduced roGFP2. Arrows indicate the time point of addition of 40 μM GSSG. As oxidation and reduction controls (calibration), 1 μM of roGFP2 was treated with 10 mM of DTT or 10 mM H_2_O_2_; n = 3 ± SDs.

As class I GRXs show oxidoreductase activity on disulfides that are formed and released via a *S-*glutathionylation intermediate (Lillig et al., 2008; Deponte, 2022), redox-sensitive GFP2 is a suitable target protein for *in vitro* assays, providing a direct fluorescent read-out for disulfide redox state (Meyer et al., 2007; Gutscher et al., 2008; Schwarzländer et al., 2008; Meyer and Dick, 2010). Hence, we tested PpGRXC5 oxidizing and reducing activity on recombinant roGFP2, in a direct comparison with the previously characterized AtGRXC1 (Trnka et al., 2020) **(Fig. 2B)**. In the *in vitro* roGFP2 reduction and oxidation assays PpGRXC5 was able to reduce and oxidise roGFP2 kinetically faster than AtGRXC1, confirming high activity in the reductive half-reaction and additionally showing high activity in the oxidative half-reaction of thiol-disulfide oxidoreductase function. Thus, PpGRXC5 is a typical class I GRX that shows efficient (de)glutathionylation as well as thiol-disulfide oxidoreductase activities.

### The single stromal class I GRX PpGRXC5 is dispensable and not a main cause for the dwarfism in **Δ***gr1* mutants

To generate null mutants of PpGRXC5 in *P. patens* we used homologous recombination **(Fig. 3A)**. We confirmed several independent transgenic lines of which we used the null mutants Δ*grxc5*#54 and Δ*grxc5*#249 **(Fig. S1)** for further experimentation. Growth under standard conditions was comparable to the wild-type **(Fig. 3B)**, which is in contrast to the previously characterised dwarfed *P. patens* glutathione reductase knockout lines (Δ*gr1*) that exhibit a less reducing stromal steady-state *E*_GSH_ (Müller-Schüssele et al., 2020). As PpGRXC5 can contribute to plastid GSSG generation or mediate *S*-glutathionylation **(Fig. 1)**, we additionally tested whether the defects observed in Δ*gr1* were dependent on GRXC5 activity. To this end, we generated Δ*gr1/* Δ*grxc5* double knock-outs by transforming the Pp*GR1* knock-out construct (Müller-Schüssele et al., 2020) into protoplasts of Δ*grxc5*#54 and confirmed the correct integration and absence of transcript for *GR1* **(Fig. S1)**. Lack of GRXC5 did not rescue the Δ*gr1* phenotype, as Δ*gr1/* Δ*grxc5* mutant lines were dwarfed **(Fig. 3B)**. Quantification of growth revealed a trend for increased fresh weight of Δ*gr1/*Δ*grxc5* compared to Δ*gr1*, that was not statistically significant **(Fig. S2)**. In addition, Δ*gr1/*Δ*grxc5* showed the same sensitivity to oxidative stress as Δ*gr1* (H_2_O_2_ treatment, **Fig. 3C)**. Thus, neither GSSG generation by PpGRXC5 nor an involvement in a putative toxic *S*-glutathionylation is contributing to the major defects observed in Δ*gr1* lines.

**Figure 3:**
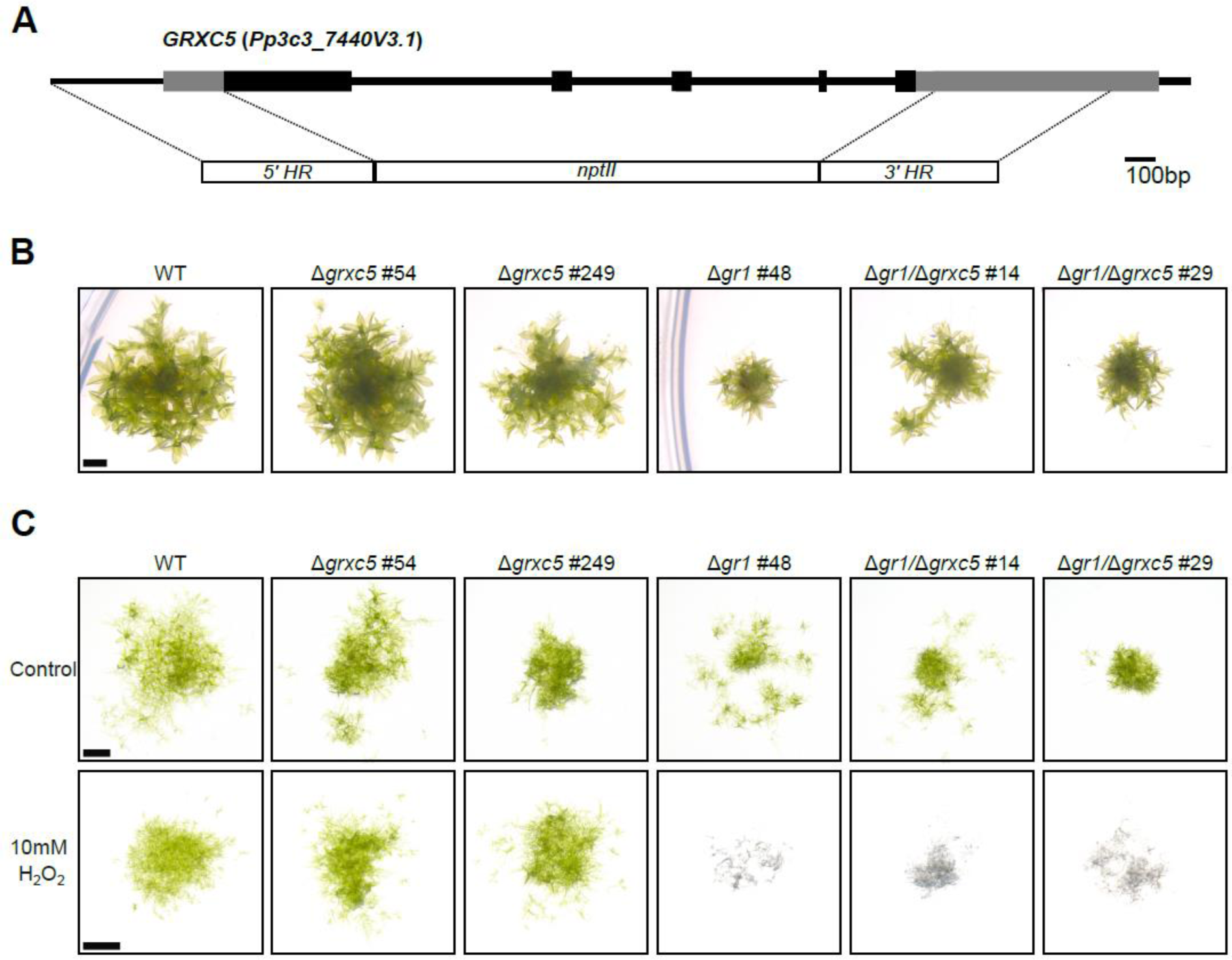
Phenotype of Δ*grxc5,* Δ*gr1* and Δ*gr1/*Δ*grxc5* plants. **(A)** Schematic overview of the Pp*GRXC5* gene structure and knock-out construct; exons = boxes; UTRs (untranslated regions) light grey, coding sequence black; HR, homologous regions; *nptII*, neomycin phosphotransferase resistance cassette. **(B)** *P. patens* grown on KNOP-ME pH 5.8 agar plates in 100 μmol photons m^-2^s^-1^ (16 h light, 8 h dark) for four weeks, scale bar = 1mm. Row represent colonies grown on the same plate. **(C)** Protonema culture spotting assay on KNOP-ME plates after incubating with 10 mM of H_2_O_2_ for 15 minutes as oxidative challenge. A 20 µl aliquot of protonema culture was placed on a KNOP-ME agar plate and grown under 60 μmol photons m^-2^s^-1^ (16 hours light, 8 hours dark) for 7 days. Control cultures were treated equally except for addition of H_2_O_2_. Images were taken after 7 days of recovery. Scale bar: 1 mm.

To further test for altered stress resilience or growth defects in Δ*grxc5* lines, we exposed plants to low light, fluctuating low light/high light and heat, but did not identify any morphological differences to WT **(Fig. S3)**. To test for differences in photosynthetic light reactions, we determined light-induction and relaxation kinetics, calculating non-photochemical quenching (NPQ) under control conditions and after exposure to high light (450 μmol photons m^-2^ s^-1^). Δ*gr1* plants showed increased non-photochemical quenching (NPQ) combined with retarded relaxation kinetics consistent with a previous report (Müller-Schüssele et al., 2020), while Δ*grxc5* lines did not show any significantly different response in NPQ induction or relaxation **(Fig. S4)**. Our results confirm that neither the growth defect nor the sensitivity to high light of Δ*gr1* plants is detectable in Δ*grxc5* plants, indicating that the negative effects of the less negative *E*_GSH_ in Δ*gr1* plants are not mediated via GRXC5. Instead, Δ*grxc5* plants showed WT-like growth in all long-term stress assays.

### Lack of class I GRX leads to distinct roGFP2 oxidation state and changed reduction kinetics after oxidative challenge in the stroma

As we did not identify phenotypical deviations of Δ*grxc5* lines from WT macroscopically, we sought to investigate stromal redox dynamics *in vivo* by introducing a genetically encoded biosensor. Redox-sensitive GFP2 (roGFP2) specifically equilibrates with the steady-state *E*_GSH_ of the local subcellular compartment (Meyer et al., 2007; Schwarzländer et al., 2008). As this equilibration is catalysed by class I GRX, human GRX1 (hGrx1) fused to roGFP2 (Grx1- roGFP2) is the standard probe used, to guarantee independence from local endogenous GRX activities (Gutscher et al., 2008). Since it was our aim to measure the endogenous GRX activity we exploited the requirement of GRX-mediated equilibration and targeted roGFP2 without fused hGRX1 to the plastid stroma of WT and Δ*grxc5#54* genetic backgrounds. We generated a construct for expression of roGFP2 fused to the transketolase targeting peptide (TKTP) (Schwarzländer et al., 2008; Speiser et al., 2018)), constitutively driven by the Pp*Actin5* promoter, targeting roGFP2 to the plastid stroma. Exclusive targeting of roGFP2 to plastids in stable transgenic lines was confirmed microscopically **(Fig. S5)** and sensor lines for each background were characterised and selected for further analyses (*TKTP-roGFP2* in Δ*grxc5#54* lines #17 and #21; *TKTP-roGFP2* in WT #20).

Sensor responsiveness to oxidation/reduction *in vivo* was confirmed via addition of either 10 mM DTT (reductant) or 5 mM DPS (2,2’-dipyridyl disulfide, thiol-specific oxidant) to moss protonema. Using a fluorescence plate-reader, excitation spectra for roGFP2 fluorescence were recorded **(Fig. 4A)**, revealing a comparable dynamic range (between 2.0 and 2.2 (405/488)) of the sensor response in WT and Δ*grxc5* background. In comparison to TKTP- roGFP2 WT#20, the physiological (untreated, Phys.) spectra of Δ*grxc5* sensor lines revealed lower excitation above the isosbestic point (c. 425 nm) and higher excitation below 425 nm, showing increased oxidation of roGFP2 of c. 20% (calculated for 405/488 nm). We additionally quantified this oxidation by ratiometric analysis of confocal images of the respective lines and found an increase in median 405/488 ratio of 1.64 ± 0.59 in Δ*grxc5#17,* 1.54 ± 0.40 in Δ*grxc5#21* compared to 1.15 ± 0.38 in WT#20 **(Fig. 4B)**. Taking into account sensor calibration (*i.e.*, ratio values for complete reduction and oxidation *in vivo*), these values support a by c. 20% higher degree of oxidation (OxD) on stromal roGFP2, compared to WT, in untreated cells.

**Figure 4:**
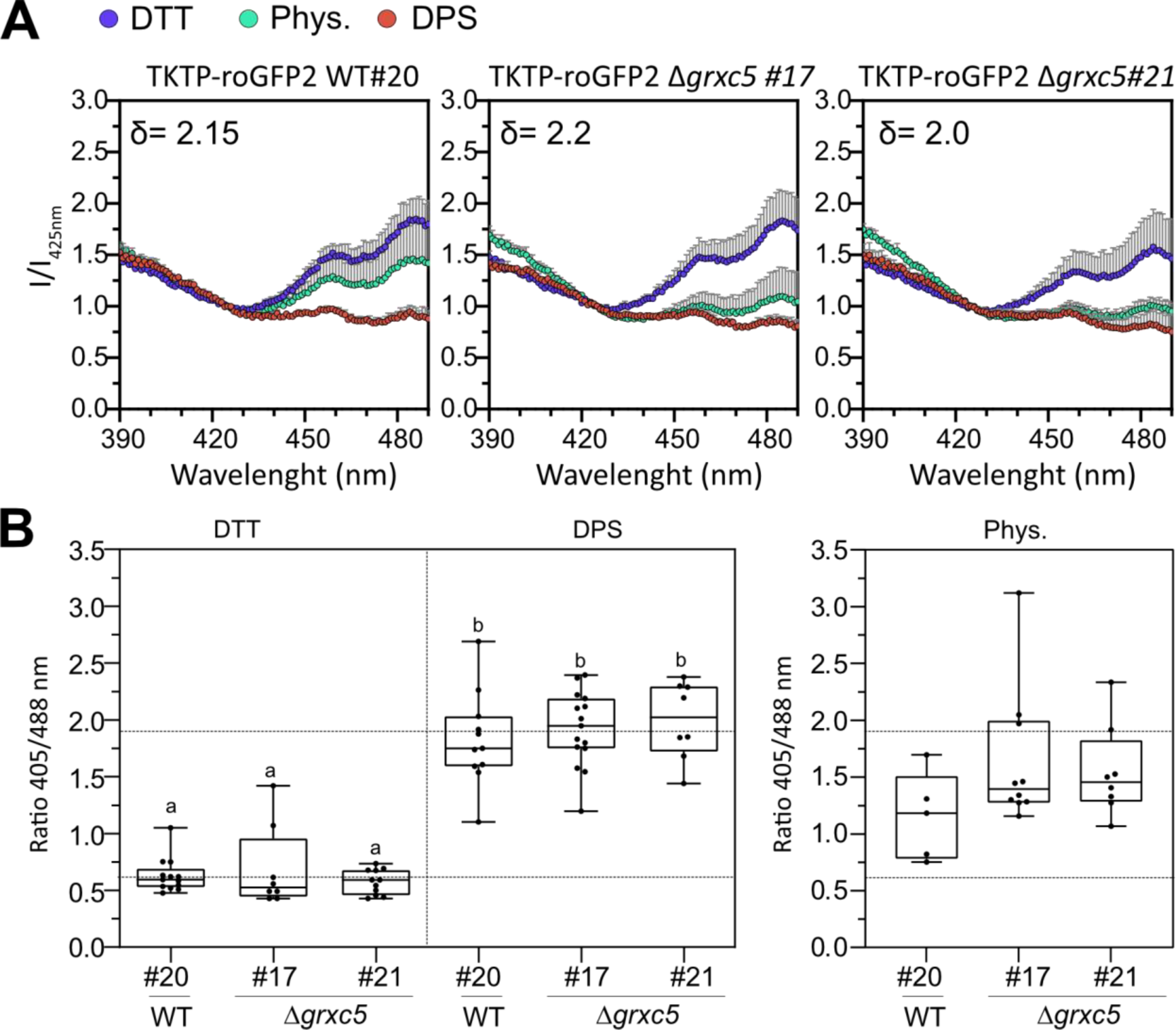
Redox state of stroma-targeted roGFP2 in WT and Δ*grxc5*. **(A)** *In vivo* excitation spectra: protonema cultures of *P. patens* were not treated (Phys.) or treated for 30 min with 10 mM DTT or 5 mM DPS. Fluorescence was excited from 390 to 490 nm while emission was collected at 535-16 nm in a plate-reader based setup. Intensities were normalized to the intensity of the isosbestic point of roGFP2 (425 nm); n=3, mean + SD. Delta depicts the dynamic range (405/488 nm) for each line. **(B)** Left panel: Image-based sensor calibration of TKTP-roGFP2 in *P. patens* with 10 mM DTT or 5 mM DPS incubated for at least 20 min before imaging with the CLSM (ex. 405, 488 nm; em. 508-535 nm); n>7, box blots: line=median and whiskers=min/max values; two-way ANOVA and Tukey’s multiple comparison test was conducted (p<0.0001), different lowercase letters indicate significant difference, dynamic range c. 3.2. Right panel: image-based analysis of steady state sensor ratio (405/488 nm) under physiological conditions: *P. patens* protonema/gametophore culture was pre-incubated in the dark for 30 min before imaging with the CLSM (ex. 405, 488 nm; em. 508-535 nm); n=5-10 pictures, box blots: line=median and whiskers=min/max values, one-way ANOVA and Tukey’s HSD test (ratio∼genetic background, p=0,08); horizontal lines: 0% and 100% oxidation based on mean DTT and mean DPS values, according to sensor calibration, see left panel.

As higher OxD values for roGFP2 are interpreted as a less negative stromal *E*_GSH_, at least in the presence of GRX activity, we additionally quantified total GSH levels by HPLC from five biological replicate samples of protonema and found a trend for an increase in total glutathione in Δ*grxc5*#54 (p=0.15) and a significant increase in total glutathione in Δ*grxc5*#249 (p=0.0006), compared to WT (one-way ANOVA, Tukey’s multiple comparison) **(Fig. S6)**. These analyses suggest that loss of PpGRXC5 causes a higher steady state OxD of roGFP2 in the stroma, as well as a trend to increased GSH levels.

As long-time exposure to various stresses did not reveal any defects in Δ*grxc5* lines, we simulated a pulse of oxidative stress to generate a temporary oxidation and compared the subsequent recovery. We exposed moss protonema to 10 mM H_2_O_2_, which causes complete oxidation of roGFP2. After removal of H_2_O_2_, we monitored the recovery kinetics of roGFP2 reduction state. While the WT background showed a fast recovery almost immediately after removal of H_2_O_2_, Δ*grxc5* null mutants showed slower, approximately linear recovery of roGFP2 redox state **(Fig. 5A)**. Using lower H_2_O_2_ concentrations in a 1-5 mM range, fast recovery was already observable during the presence of exogenously added oxidant in the WT background while Δ*grxc5* null mutants consistently showed slower, approximately linear recovery from oxidative stress **(Fig. S7)**. To additionally monitor oxidation kinetics of the roGFP2 disulfide, we pre-reduced moss protonema and subsequently used injection of H_2_O_2_ combined with 1.5 s measuring intervals. We found that oxidation of roGFP2 in Δ*grxc5* stroma proceeded equally fast or slightly faster **(Fig. 5B)**. Thus, disulfide formation in response to severe oxidative stress is fast in Δ*grxc5* null mutants while stromal GSH-dependent disulfide reduction is slow and almost linear.

**Figure 5:**
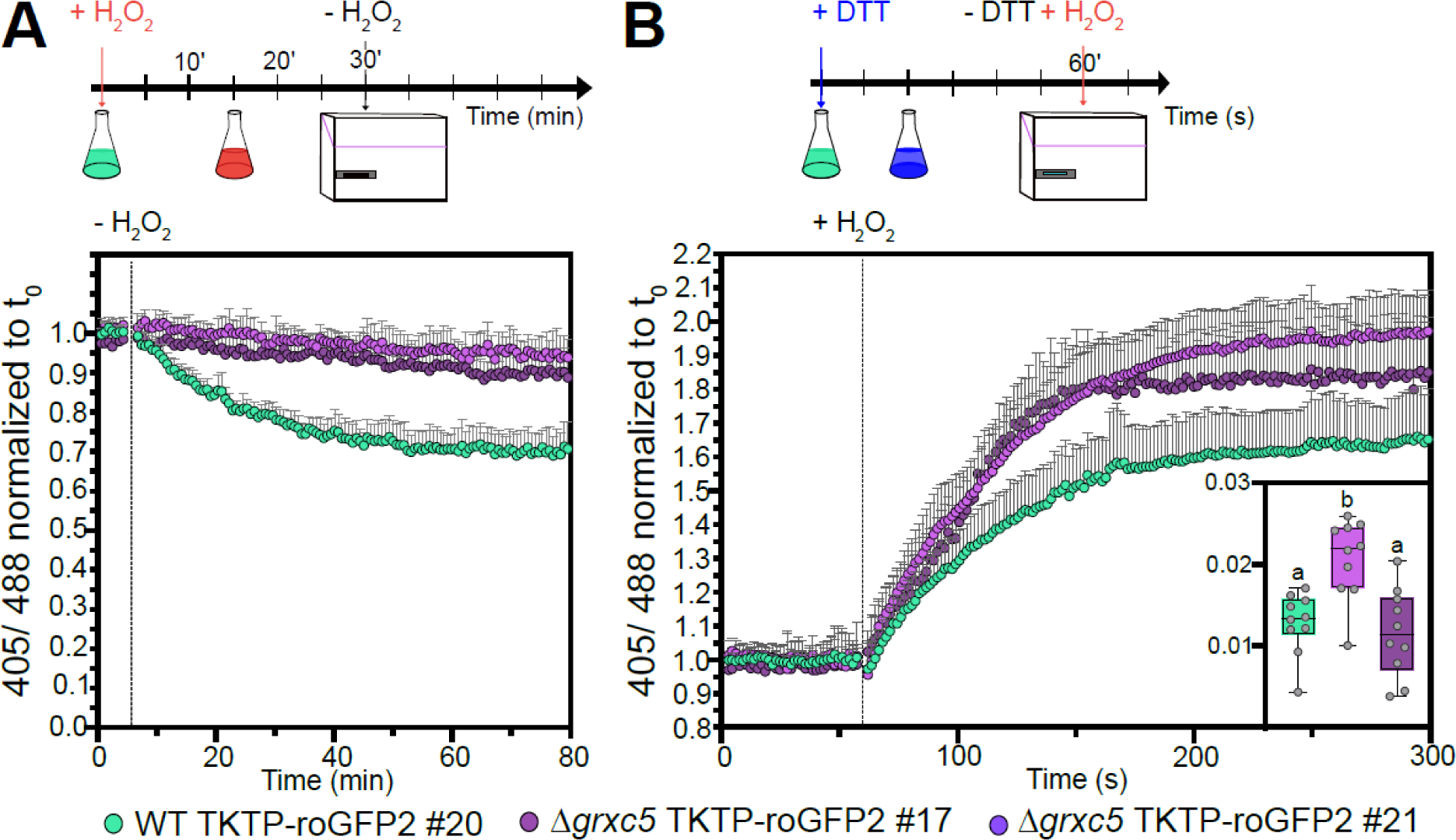
*In vivo* kinetics of stromal roGFP2 in response to oxidative challenge. **(A)** Protonema culture expressing *TKTP-roGFP2* was treated with 10 mM H_2_O_2_ for 30 min and H_2_O_2_ subsequently removed and exchanged with imaging buffer to monitor recovery from oxidative challenge. 405/488 nm ratio of n=6 biological replicates was normalized to t_0_, graph depicts mean+SD. A two-way ANOVA with Tukey’s multiple comparison test was conducted for each time point revealing significant differences in roGFP2 kinetics between WT and both Δgrxc5 #17 and Δgrxc5#21 starting from 11 min after peroxide removal (p< 0.001, n=6). **(B)** *In vivo* oxidation rates after injection of H_2_O_2_ (final concentration = 2 mM) and monitoring of oxidation rate every 1.55 s. Protonema samples were pre-reduced using 10 mM DTT. Shown are the mean + SD, n=10. Slope (inset, ΔR/Δt) was calculated for the first 10 s after injection (eight data points). Box plot whiskers depict min and max values with the horizontal line indicating the median. One-way ANOVA with Tukey’s multiple comparison test was conducted to test for significant difference (p<0.027).

**Figure 6.**
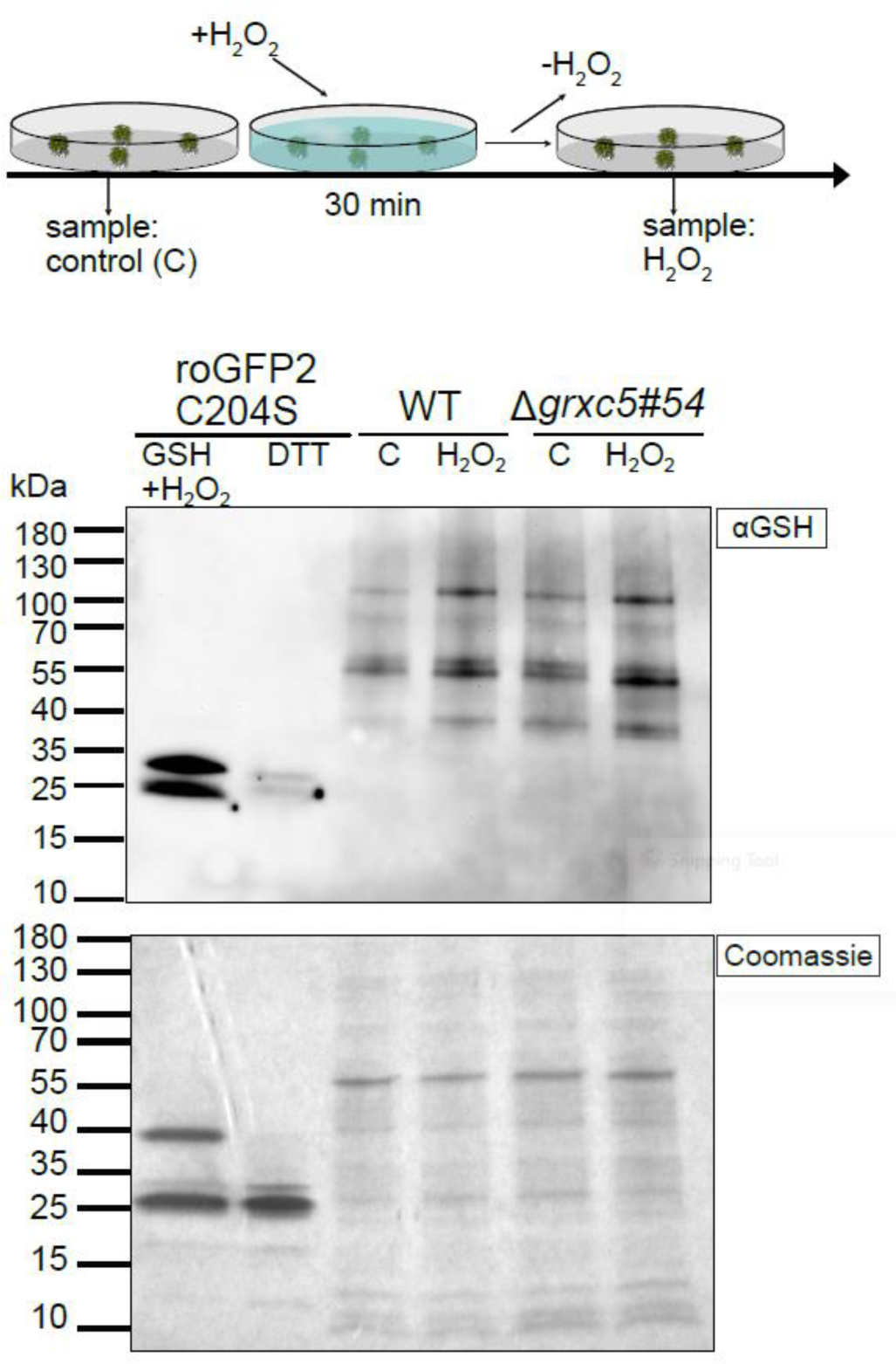
Total protein-bound glutathione after oxidative challenge. **(A)** Schematic overview of the experimental set-up and sampling. **(B)** Immunoblot using α-GSH (ThermoFisher). Total protein extracts from *P. patens* WT and Δ*grxc5*#54 gametophore tissue either non-treated (‘C’) or treated with 10 mM H_2_O_2_ for 30 min (‘H_2_O_2_’) (see panel A); cysteine oxidation was blocked with 20 mM NEM (*N*-ethyl maleimide) in the lysis buffer. As loading control, 10 µg total protein was loaded onto a 4-20% gradient non-reducing SDS-PAGE lower panel). As control for antibody specificity, purified roGFP2C204S (10 µM) was treated with 10 mM H_2_O_2_ in the presence of 2 mM of GSH for 30 min (positive control, glutathionylated roGFP2C204S) or treated with 10 mM DTT (negative control, no glutathionylated roGFP2C204S), and 12 µl loaded per lane.

### Linking *S*-glutathionylation levels to oxidative challenge and GRX activities

Using roGFP2 enabled us to observe disulfide formation and reduction kinetics in the absence of a class I GRX under different environmental conditions. However, disulfides would only be formed in response to *S*-glutathionylation if a second cysteine is present in a suitable distance to reduce the mixed disulfide, forming an intramolecular disulfide and concomitant release of GSH **(Fig. 1)**. While this is the case for roGFP2, many *in vivo* targets of class I GRXs may remain in a *S*-glutathionylated state (Zaffagnini et al., 2012b; Müller-Schüssele et al., 2021a). Hence, we investigated if total protein *S*-glutathionylation levels are altered in Δ*grxc5* null mutants, using Western blotting with an anti-GSH antibody **(Fig. 6)**. As a control, we incubated *in vitro* purified roGFP2 mutant lacking the resolving Cys204 (roGFP2C204S) (Trnka et al., 2020) with 10 mM H_2_O_2_ in the presence of 2 mM GSH, inducing *S*-glutathionylation. Protein extracts from non-stressed and stressed WT and Δ*grxc5* plants both showed equally increased signal after immunodetection of protein-bound GSH, starting from a similar level. Increase of protein-bound GSH was consistent and independent of presence of GRXC5 after oxidative challenge **(Fig. S8)**.

### Stromal GSH-dependent redox kinetics are altered in dark/light/dark transitions, without an effect on carbon fixation

As stromal Grx1-roGFP2 showed a previously unknown light/dark transition-dependent *E*_GSH_ dynamics in WT (Müller-Schüssele et al., 2020), we next assessed stromal roGFP2 redox dynamics in Δ*grxc5* null mutants using both a plate-reader based setup **(Fig. 7A)** and a confocal microscopy-based setup (Müller-Schüssele et al., 2020) **(Fig. 7B)**. We found that light/dark transition-dependent dynamics of stromal roGFP2 oxidation state were also observable using roGFP2 without fused Grx1 in WT background: a sudden light to dark transition leads to rapid sensor oxidation, followed by a recovery phase. In contrast, light/dark dependent roGFP2 redox changes were completely absent in Δ*grxc5* null mutants. Responsiveness of the roGFP2 probe was confirmed after light/dark treatment by calibration **(Fig. 7A)**, demonstrating that roGFP2 redox state in Δ*grxc5* was well inside dynamic range of roGFP2. As a complementary approach, we used a CLSM-based setup where we were also able to follow roGFP2 redox state in the light and confirmed absence of dark/light/dark transition-dependent roGFP2 redox dynamics in Δ*grxc5* **(Fig. 7B)**.

**Figure 7:**
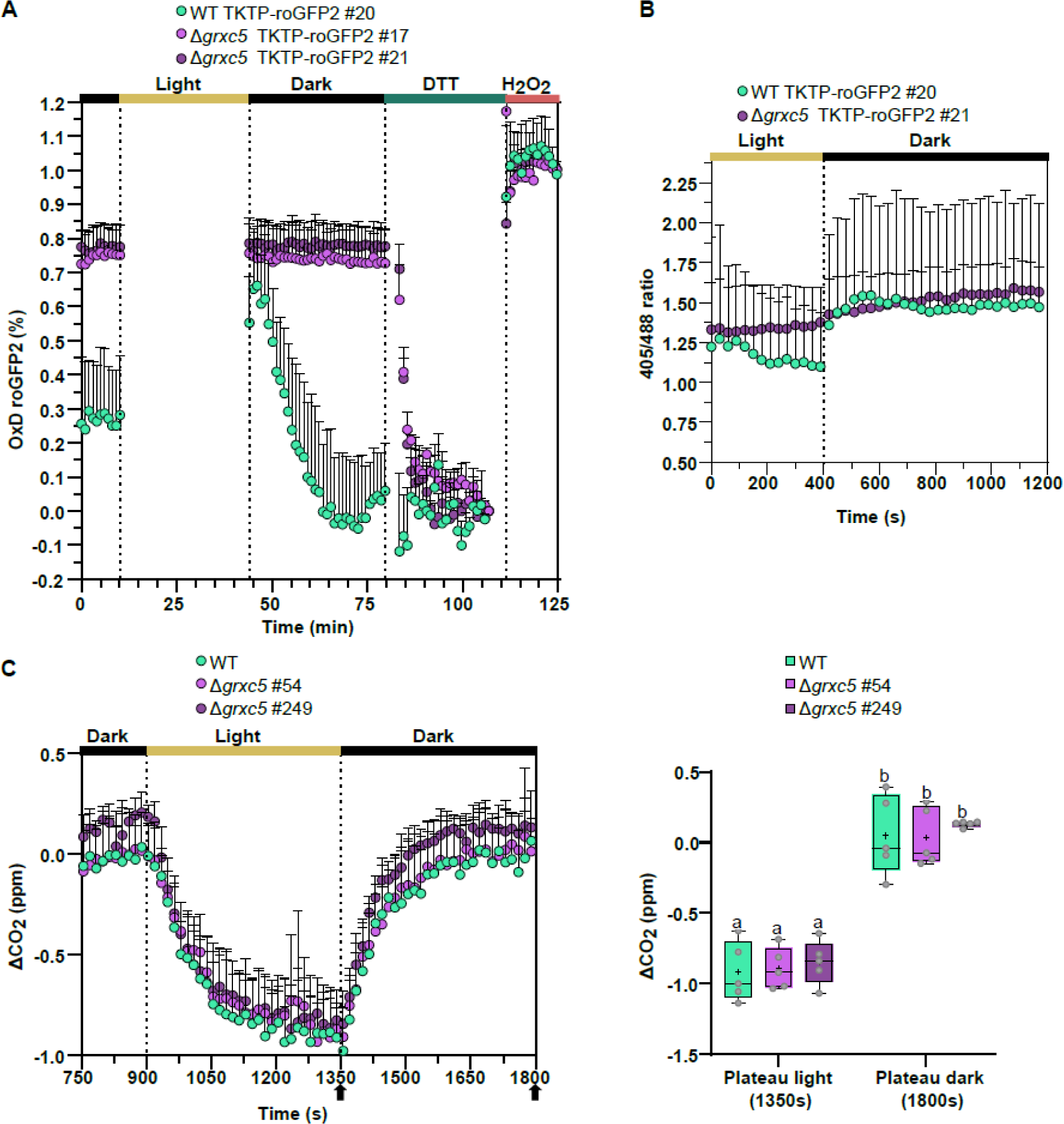
Light-dependent roGFP2 dynamics and CO_2_ assimilation during dark-light-dark transitions in protonema culture of *Δgrxc5* and WT. **(A)** Reduction/oxidation dynamics in dark/light/dark transitions. In a 96-well plate with 200 µl of protonema culture from *P. patens*, initial fluorescence was measured after 30 min dark incubation using a plate reader-based setup. Subsequently, the plate was illuminated for 30 minutes to an intensity of ∼200 µmol photons m^-2^s^-1^ using external LED illumination. After dark/light/dark transition, each well was calibrated by first replacing the buffer with 10 mM DTT and then10 mM H_2_O_2_. OxD = degree of oxidation, shown are the mean (+SD) of n=3. **(B)** Image-based analysis of oxidation/reduction dynamics of TKTP-roGFP2 in *P. patens* gametophores and protonema tissue grown in liquid culture. CLSM time series of dark-adapted samples: 1 min in the dark, illumination by external light source for 5 min (100 µmol photons m^-2^s^-1^), followed by dark incubation. Images were taken every 30 seconds for 20 minutes; shown is the mean + SD of n=7-8. **(C)** Left panel: changes in CO_2_ partial pressure during dark/light/dark transitions in protonema culture of Δ*grxc5* lines and WT using a 7.5 min light and 7.5 min dark cycle (98% humidity, 500 ppm CO_2_, 22 °C, and 75 µmol photons m^-2^s^-1^). The absolute changes in CO_2_ levels were measured after zero-point (ZP) subtraction (nylon membrane filter wetted with KNOP-ME as ZP). Right panel: Changes in CO_2_ partial pressure after reaching plateau values during the light and dark phases, respectively (indicated by arrows in left panel). One-way ANOVA and Tukey’s multiple comparison to assess significant differences between Δ*grxc5* lines and WT at the end of the light cycle (1350s) and the end of the dark cycle (1800s): p=0.99. Boxes display the 25-75 percentiles, with the minimum and maximum values indicated by the whiskers, and the median marked by the horizontal line (n=5).

As our results suggest potential differences in glutathione-dependent redox dynamics in the chloroplast stroma, we assessed cross-talk to the TRX system by quantifying CO_2_ release and assimilation from protonema in a dark/light/dark transition **(Fig. 7C)**. Our analysis did not find any significant differences in either CO_2_ release in the dark, nor in CO_2_ uptake in the light, suggesting overall unaltered redox-regulation of the CBB cycle enzymes in ***Δ****grxc5* lines, compared to WT. Thus, fast responses of the roGFP2 redox state to light/dark dependent *E*_GSH_ dynamics are absent in ***Δ****grxc5* plants, while regulation of carbon fixation was unaltered under the tested conditions.

## Discussion

### Glutathione as an electron donor in the plastid stroma

Thiol redox states depend on their reaction kinetics with different small molecules or protein reductants and oxidants, including thiols in a suitable distance for disulfide formation. Different thiol/oxidised thiol redox couples can be in thermodynamic equilibrium after incubation in the range of minutes to hours, whereas enzymatically catalysed equilibration takes place in the range of seconds to minutes. GRX-catalysed redox equilibration depends on GSH as electron donor. We determined the enzyme kinetics of PpGRXC5 (YCPYC active site) *in vitro* and found high deglutathionylation activity (HED assays) as well as thiol-disulfide oxidoreductase activity (roGFP2 *in vitro* assays). The apparent second-order rate constant(s) (*k*_cat_^app^/*K*_m_^app^) for PpGRXC5-catalyzed reactions (HED assay) was similar to AtGRXC1 (YCGYC active site) (Riondet et al., 2012) or AtGRXC5 (WCSYC active site) (Couturier et al., 2011), showing that GRXC5 is an evolutionary conserved redoxin (Müller-Schüssele et al., 2021a) with a typical class I GRX functionality as GSH-dependent (de)glutathionylation and thiol/disulfide oxidoreductase activity.

Disulfides can be characterized by their midpoint potential, *i.e.*, the redox potential at which 50% of the molecules are reduced and 50% oxidized. The disulfide formed by the genetically encoded biosensor roGFP2 is well-characterised, with a consensus midpoint potential of - 280 mV (pH 7), and enzymatically catalysed by GRX in dependence on *E*_GSH_ ((Dooley et al., 2004; Meyer et al., 2007; Gutscher et al., 2008; reviewed in Meyer and Dick (2010); Schwarzländer et al. (2016); Müller-Schüssele et al. (2021b)). We targeted this redox sensor protein to the plastid stroma, without the fusion to a GRX, to track local GSH-coupled redox dynamics in the model moss *P. patens*. In the stroma of WT plants, roGFP2 redox state indicated a steady-state stromal *E*_GSH_ of c. -312 mV (pH 8), consistent with the -311 mV (pH 8) measured using Grx1-roGFP2 in WT background of *P. patens* plastid stroma (Müller-Schüssele et al., 2020). A similar physiological steady-state stromal *E*_GSH_ in the dark-adapted state of WT using either roGFP2 or Grx1-roGFP2 confirms similar functionality of roGFP2 in WT stroma, without fused hGrx1. Compared to WT, we detected an increased 405/488 nm ratio and corresponding increased roGFP2 degree of oxidation (OxD) of c. 20% in the stroma of Δ*grxc5* lines. In addition, we found a trend for increased total GSH content, which would according to the Nernst equation suggest more negative *E*_GSH_, although HPLC measurements do not allow for subcellular resolution. A shift of roGFP2 thiol/disulfide redox state to more oxidized values usually reveals an oxidative shift in *E*_GSH_, as roGFP2 specificity for the glutathione/GRX system was tested *in vitro* and *in vivo* (Meyer et al., 2007; Begas et al., 2017; Trnka et al., 2020; Schlößer et al., 2023). Thus, inefficient reduction of GSSG caused by absence of glutathione reductase (GR) leads to an increase of OxD_roGFP2_ in the same subcellular compartment (Marty et al., 2009; Marty et al., 2019; Müller-Schüssele et al., 2020).

In Δ*gr1* lines, a shift of c. 44% in OxD_roGFP2_ was measured using stroma-targeted Grx1-roGFP2 which reports an oxidative stromal *E*_GSH_ shift of c. 33 mV (Müller-Schüssele et al., 2020). In theory, the measured shift in roGFP2 redox status in the Δ*grxc5* lines may either indicate altered *E*_GSH_, *i.e.*, local decrease in GSH and/or local increased GSSG concentration, or inefficient equilibration between *E*_GSH_ and roGFP2 (or a combination of both). The following argument supports an oxidative shift in OxD_roGFP2_ in the Δ*grxc5* stroma independent of a change in local *E*_GSH_: (1) GR is still present in the stroma of Δ*grxc5*, safeguarding the highly reducing stromal *E*_GSH_. (2) *In vivo* stromal roGFP2 reduction rates after oxidative challenge (**Fig. 5A**) are similar (i.e. in the range of minutes to hours) to thermodynamically driven *in vitro* reduction rates based on highly negative *E*_GSH_, but lacking addition of GRX (**Fig. 2B**, (Meyer et al., 2007)). Thus, without class I GRX present, a shifted OxD_roGFP2_ likely originates from a decreased glutathione-dependent reduction rate of the roGFP2 disulfide, never reaching thermodynamic equilibrium with *E*_GSH_. Instead the interaction with other redox couples becomes apparent in the form of the oxidation (i.e. unchanged oxidation rates), resulting in an OxD_roGFP2_ steady state further away from thermodynamic equilibrium with glutathione as electron donor (schematic model **Fig. 8**). After oxidative challenge, lacking GRXC5 activity leads to slow recovery rates from higher thiol oxidation levels (**Fig. 8**). Thus, Δ*grxc5* and Δ*gr1* mutants both show altered OxD_roGFP2_ *in vivo*, but through different mechanisms, *i.e.*, lacking reduction of GSSG *vs*. kinetic uncoupling between *E*_GSH_ and disulfide redox state. Our data show that fast enzymatically catalysed equilibration of roGFP2 redox state with the highly reduced glutathione pool is not complemented by other redox systems present in the stroma and confirm PpGRXC5 as the sole stroma-targeted class I GRX in *P. patens*.

**Fig. 8:**
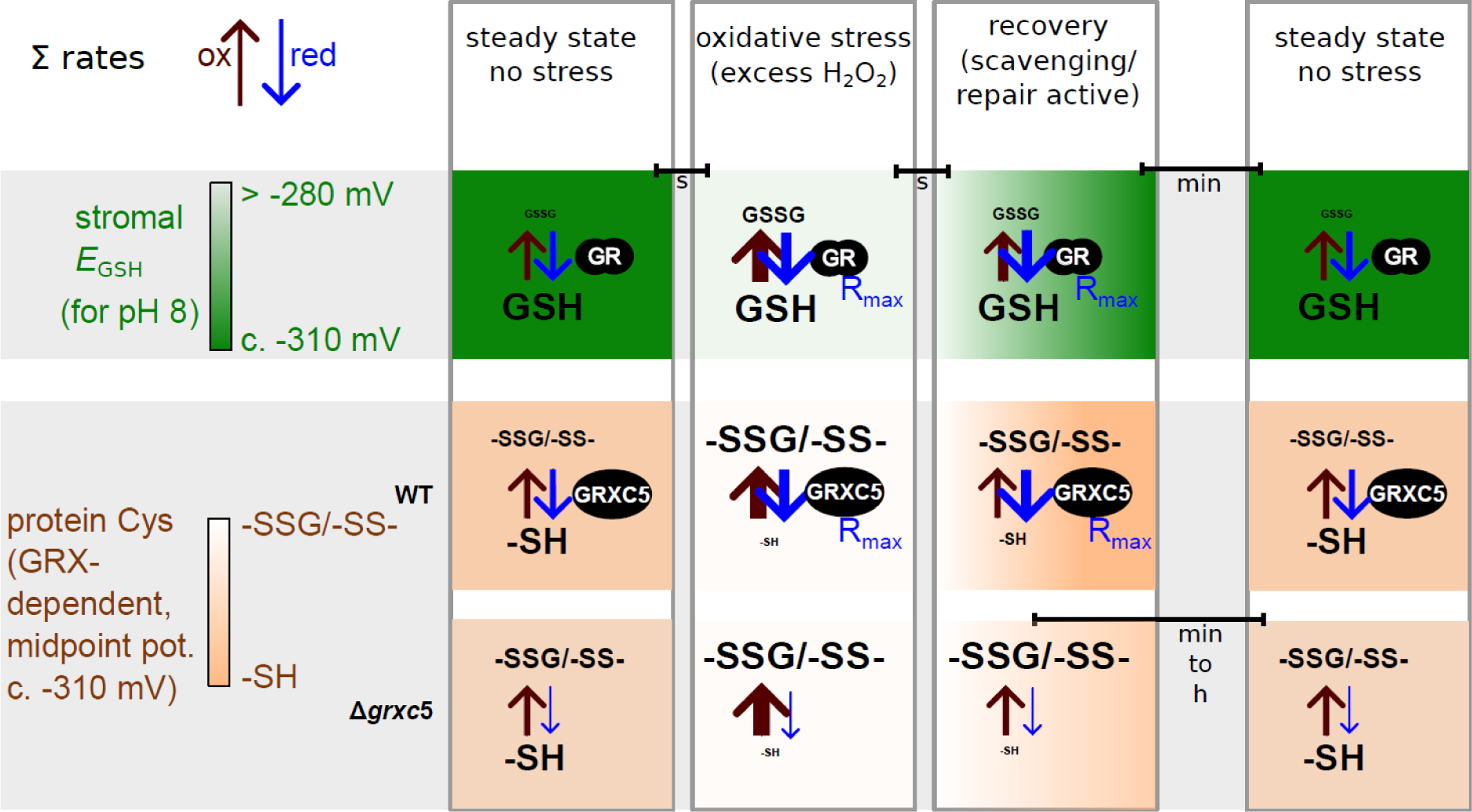
Schematic overview: Model of observed dynamic redox states as a consequence of changing oxidation and reduction rates. Simplified scheme of steady states of the GSH/GSSG redox couple and resulting *E*_GSH_, as well as the protein thiol/disulfide and thiol/glutathionylated redox couples before, during and after an oxidative challenge, as investigated in this work using roGFP2. Redox potentials are given for pH 8 (midpoint potential of roGFP2 at pH 8 is -310 mV). Colour scales exemplify relative changes between reduced and oxidised forms. Bars indicate approximate durations of transitions. Arrow thickness indicates sum of oxidation rates (i.e. all direct oxidation (slow) and enzymatically catalysed oxidation (fast)) and sum of reduction rates that would result in the observed profile of redox dynamics. Lack of GRXC5 leads to more oxidised steady state for target cysteines, as well as slow recovery rates. Enzymes are depicted as black ovals. R_max_ = maximal reachable reduction rate with current enzyme copy number.

Moreover, by generating Δ*grxc5/*Δ*gr1* double mutant lines, we found that GRXC5 function is not a main cause for the stress-sensitive and dwarfed Δ*gr1* phenotype. This excludes GRXC5 as main GSSG-producing enzyme in the stroma, as well as putatively toxic GRX-dependent *S*-glutathionylations occurring in response to an oxidatively shifted *E*_GSH_. Glutathione can serve as electron donor for additional stromal enzymes involved in ROS-induced damage repair, signalling or ROS scavenging, such as plastidial MSRB1 (1 Cys methionine sulfoxide reductase B1) (Laugier et al., 2009), PRXIIE (Gama et al., 2008), GST iota/lambda (GSH *S*- transferase) and DHAR (dehydroascorbate reductase) (reviewed in (Müller-Schüssele et al., 2021a). These processes represent additional candidates for stroma-localised GSSG generation (see GSH oxidation rates in **Fig. 8**), and merit further investigation identifying their specific contributions to stress tolerance.

### Is class I GRX function relevant in the chloroplast stroma?

TRX and GRX functions are partially redundant in *E. coli*, mostly related to their function as alternative electron donors to ribonucleotide reductase, powering cell growth (Fernandes and Holmgren, 2004). In the cytosol of *S. cerevisiae*, Δ*grx1*/Δ*grx2* null mutants grow normally but show increased sensitivity to oxidative challenge (Luikenhuis et al., 1998; Eckers et al., 2009). Lethality only occurred when concomitantly knocking-out all cytosolic TRX isoforms (Draculic et al., 2000). GRXs can operate through a monothiol mechanism or a dithiol mechanism in which only the N-terminal active site cysteine or both active site cysteines participate in the redox process, respectively (Lillig and Berndt, 2013). One cytosolic monothiol GRX was able and sufficient to compensate for loss of all TRX and GRX (Zimmermann et al., 2020). Partial redundancy between the GSH/GRX and TRX redox systems was also demonstrated in plants, regarding the cytosol and mitochondrial matrix (Marty et al., 2009), but not for the plastid stroma (Marty et al., 2019). Regarding mutants lacking all class I GRX of one compartment, double null mutants of *A. thaliana* lacking cytosolic GRXC1 and GRXC2 were originally described to be lethal (Riondet et al., 2012) but have recently been reported to be viable and with a WT-like phenotype (Schlößer et al., 2023). In the secretory pathway, which lacks a GR and thus possesses a less reducing *E*_GSH_, double mutants of GRXC3/GRXC4 grow normally (Schlößer et al., 2023) but showed decreased hypocotyl length compared to WT when grown in elevated temperature (Dard et al., 2022). Regarding plastid-targeted class I GRX, no null mutants were described yet. PpGRXC5 has a CPYC active site and could operate via the monothiol or the dithiol mechanism (Deponte, 2017; Zimmermann et al., 2020). During land plant evolution, a second isoform of plastid-targeted class I GRX lacking the second active site Cys, GRXS12, evolved within the same clade (Müller-Schüssele et al., 2021a). Poplar GRXS12 was characterized as functional monothiol class I GRX (Couturier et al., 2009; Zaffagnini et al., 2012a), but the biological relevance of either class I GRX plastid isoform remains so far unclear.

While *E*_GSH_ is usually robust, except under extreme abiotic stress conditions challenging GR capacity (Wagner et al., 2019; Ugalde et al., 2021; Bohle et al., 2022) (**Fig. 8**), recent results have shown that stromal *E*_GSH_ is dynamic in response to physiological light/dark transitions (Müller-Schüssele et al., 2020; Haber et al., 2021). We found that in Δ*grxc5* stroma, these light- dependent *E*_GSH_ dynamics are not transferred anymore to glutathione-dependent disulfides, such as in roGFP2 (**Fig. 7A**), raising the question which endogenous target disulfides may respond differently (**Fig. 8**). As an obvious target for interference between the TRX and GSH/GRX system in the stroma, we tested CO_2_ release and uptake in WT compared to Δ*grxc5* lines and found no significant differences under the tested conditions (**Fig. 7B**). In accordance with the normal plant growth observed in Δ*grxc5* lines, this result indicates that TRX-dependent redox regulation of the CBB cycle (TRX-*m* and TRX-*f* isoforms) is not affected. This confirms limited cross-talk between the TRX and class I GRX in the plastid stroma.

The remaining question is why GSH/GRX and TRX-dependent redox cascades can remain largely separated in the same subcellular compartment. One possible explanation is substrate specificity, e.g. mediated by electrostatic complementarity between redoxins and their target proteins (Bodnar et al., 2023). As at least one stromal class I GRX isoform (GRXC5 or GRXS12) is evolutionarily strictly conserved, an important function is likely, but may not consist in backing-up TRX-dependent redox cascades. In accordance, plastid-localised PRXIIE that is efficiently reduced via class I GRX, does not function as TRX oxidase like other PRX (Telman et al., 2020).

### *S-*glutathionylation: A needle in the haystack or important PTM?

A remaining important question is for which plastid processes fast GRX-mediated reduction kinetics of either protein *S*-glutathionylation or GSH-dependent protein disulfides would matter. By exogenously challenging plants with H_2_O_2_, we found that kinetically fast roGFP2 reduction was absent in the stroma. This leads to prolonged disulfide persistence in a time frame of at least 30 min after such an oxidative challenge, as well as to an altered steady state OxD. Thermodynamic equilibration of purified proteins with *E*_GSH_ can take hours (Meyer et al., 2007). In contrast, stromal roGFP2 oxidation kinetics after addition of H_2_O_2_ were not slower **(Fig. 5B)**. Disulfides can be directly induced by H_2_O_2_, with the rate constant being dependent on the p*K*_a_ of the more reactive cysteine (Zaffagnini et al., 2019a). In this case, the thiolate anion reacts with H_2_O_2_, forming a sulfenic acid (-SOH) and water. This sulfenic acid can react with a nearby thiol, forming a disulfide (and water). In the presence of GSH, this reaction sequence can lead to GRX-independent *S*-glutathionylation. Using an anti-GSH antibody on stressed and control samples of WT and Δ*grxc5* lines we found increased total *S*-glutathionylation after oxidative challenge. Lack of GRXC5 did not interfere with the level of glutathionylated proteins after oxidative stress treatment (**Fig. 6, Fig. S8**), indicating a minor contribution of GRXC5-mediated *S*-glutathionylation under the tested conditions. Our results support the hypothesis that the most likely scenario for protein *S-*glutathionylation (and *S*-glutathionylation-dependent disulfide formation) *in vivo* involves activated thiol derivatives such as sulfenic acids, most efficiently formed on cysteine residues that are in a deprotonated state (*i.e.*, thiolates) at physiological pH (Zaffagnini et al., 2012b).

Previous studies have revealed around 150 stromal proteins as potential *S*-glutathionylation targets, using different experimental approaches (reviewed in Zaffagnini et al. (2019a) and Müller-Schüssele et al. (2021a)). However, it is still unclear under which physiological conditions which fraction of principally susceptible cysteine residues really is *S*- glutathionylated *in vivo*, including consequences to activity, stability or localization of the affected protein molecules. If *S*-glutathionylation only occurs for a minor fraction of proteins or a minor fraction of protein molecules of one protein isoform, effects on metabolic fluxes would be unlikely, as there are still many non-glutathionylated protein molecules present in the entire population. However, protein cysteines with stable thiolate anions are interesting candidates to sense H_2_O_2_ or GSSG levels in signalling processes, with reduction rates mediated via class I GRX. In this regard, cytosolic GAPDH provides an interesting example. This enzyme contains a reactive cysteine which is essential for catalysis and is a major target of H_2_O_2_-dependent oxidation (Trost et al., 2017; Talwar et al., 2023). The subsequent reaction of GSH with the sulfenylated catalytic cysteine induces *S*-glutathionylation and protects the enzyme from irreversible oxidation. However, the persistence of the glutathionylated state, which causes an unavoidable loss of function, has a drastic and irreversible effect on structural stability inducing protein misfolding and aggregation (Zaffagnini et al., 2019b).

In order to understand the biological relevance of protein *S*-glutathionylation as PTM in the chloroplast stroma, identification of *in vivo* targets of GRXC5 (and/or GRXS12 in angiosperms) is necessary. This task will require fitting methodological tools for high-throughput protein redox state analysis in combination with suitable mutants and time frames. Life imaging with genetically encoded redox sensors can meet the challenge to follow oxidation and reduction rates *in vivo*, and to effectively dose stress treatments.

Based on our results, we conclude that stromal class I GRX are necessary to quickly release *S*-glutathionylation or disulfides formed via an *S*-glutathionylation intermediate. The question of why GRX-assisted glutathione-dependent catalysis evolved (Deponte, 2013; Deponte, 2022) is still open, especially regarding the plastid stroma. The main difference of mutants lacking class I GRX in comparison to mutants lacking GR may be that there is still sufficiently high GSH (and sufficiently low GSSG) for GSH-dependent (GRX-independent) reduction. In absence of class I GRX activity, reduction of disulfides still occurs, but at lower rates driven by thermodynamics (**Fig. 8**).

In conclusion, the most likely class I GRX functions remain thiol protection and enzyme regulation in response to oxidative challenge (Fernandes and Holmgren, 2004; Berndt et al., 2014). Potentially, fast kinetic equilibration with *E*_GSH_ is just relevant for enzymes with an *S-* glutathionylation intermediate on catalytic cysteines, as they would be temporarily “locked” in their disulfide or -SSG form in absence of a class I GRX. In the stroma, 1Cys MSRB1 and PRXIIE are interesting candidates. Alternatively, protein Cys with low p*K*_a_ could have evolved on proteins involved in (moonlighting) signalling functions in response to oxidative challenge, which still await identification as specific GRX interaction partners.

## Competing Interest

The authors declare that they have no conflicts of interest.

## Author Contributions

FB, MZ, PT, AJM and SJM-S designed the research. FB, JR, SST, HJ, FR, AB, SB, OT, MS, SK performed experiments and analysed data. SJM-S, MSchw., AJM, PJ, MD, EN, MZ, PT supervised the research and provided resources.

FB and SJMS wrote the manuscript with contributions from all authors. All authors approved the manuscript before submission.

## Data Availability

All data needed to evaluate the conclusions in the paper are present in the paper and/or the Supporting Information.

## Acknowledgements

We thank Andreas Werle-Rutter (Molecular Botany RPTU), Maria Homagk (Chemical Signalling, INRES, University of Bonn) and Bastian Welter (University of Cologne) for technical assistance, as well as Prof. Frank Hochholdinger for support. We thank Dr. Marlene Elsässer and Dr. Mareike Schallenberg-Rüdinger for generating a Gateway version of the *PTA2_Act5* expression vector.

This work was supported by the DFG-funded Research Training Group GRK2064: ‘Water use efficiency and drought stress responses: From Arabidopsis to Barley’ (AJM, MSchw., FB and SJM-S) and via the Joint Mobility Program between the DAAD (PPP Italy 57397466) and the MIUR (Prog. n. 34433) (AJM and PT). JR was supported by a PhD grant from the University of Bologna (PhD programs in Cellular Molecular Biology). Research in SK’s laboratory if funded by DFG under Germany’s Excellence Strategy – EXC 2048/1 – project 390686111. SJM-S and OT are grateful for funding obtained from BioComp 3.0 ‘Dynamic Membrane Processes in Biological Systems’. HJ, ST and SJM-S are grateful for support by the DFG- funded Research Training Group GRK2737 ‘STRESSistance’.

## Supporting Information

**Fig. S1: Construct design and knock-out validation**

**Fig. S2: Fresh weight analysis for Δ*grxc5*, Δ*gr1* and Δ*grxc5/*Δ*gr1***

**Fig. S3: Stress phenotyping of P.patens protonema culture under fluctuating light and heat stress**

**Fig. S4: Non-photochemical quenching (NPQ) measurements of 4-week-old gametophores under low and high light regimes**

**Fig. S5: Targeting of roGFP2 to plastids**

**Fig. S6: Quantification of total GSH in GRXC5 mutant lines by HPLC**

**Fig. S7: roGFP2 oxidation state and recovery in response to externally added H_2_O_2_**

**Fig. S8: Increase of protein-bound glutathione after H_2_O_2_ treatment**

**Table S1: Primer sequences**

### Supplementary Material and Methods

**Measurement of photosynthetic parameters (light induction and relaxation curves) Glutathione measurement by HPLC**

## Notes

### Competing Interest Statement

The authors have declared no competing interest.

