## Supplementary Material for "Chloroplasts lacking class I glutaredoxins are functional but show a delayed recovery of protein cysteinyl redox state after oxidative challenge"

**Bohle et al.**

### Supplementary Figures

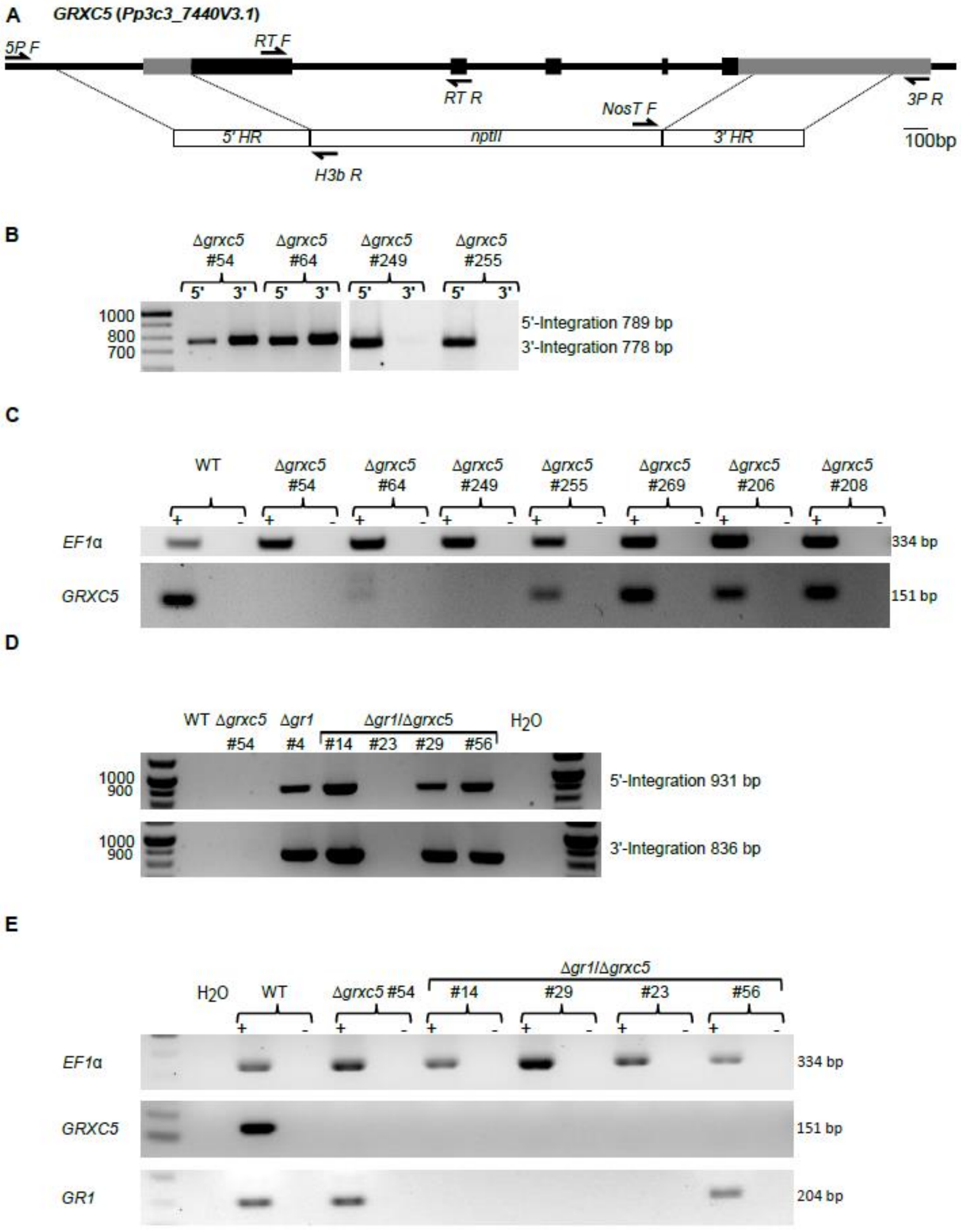


#### Supplemental Figure 1: Construct design and knock-out validation

**(A)** Schematic overview of the construct including an antibiotic resistance cassette (*nptII*) transformed via homologous recombination into the Pp3c3_7440V3 locus of *P. patens*. Exons are depicted as light grey box. Dark grey boxes display 5' and 3' UTR, arrows indicate primers used for knock-out confirmation (**Table S1**). **(B)** Testing for 5' and 3' integration of *GRXC5* knock-out construct verified by PCR using 5P_F and H3b_R (integration 5' homologous region) or NosT_F and 3P_R (integration 3' homologous region). **(C)** Verification of the absence of transcript on cDNA-level using gene-specific primers for the *GRXC5* locus (PpGrxC5ko_RT_F and PpGrxC5ko_RT_R; **Table S1**). **(D)** Testing for 5' and 3' integration of the *GR1* knock-out construct was verified by PCR using GR1_5P_F and H3b_R primers (for the integration 5' homologous region) or NosT_F and GR1_3P_R primers (for the integration 3' homologous region) (Müller-Schüssele *et al.,* 2020, **Table S1**). **(E)** Verification of the absence of *GR1* transcript on cDNA-level using gene specific primers for the *GR1* locus (PpGR1_RT_F and PpGR1_RT_R; Müller-Schüssele *et al.,* 2020). The EF1-alpha locus was used as cDNA positive control (PpEF1a_RT_F and PpEF1a_RT_R; **Table S1**). Samples without reverse transcriptase enzyme (negative control for cDNA) are indicated as '-'.

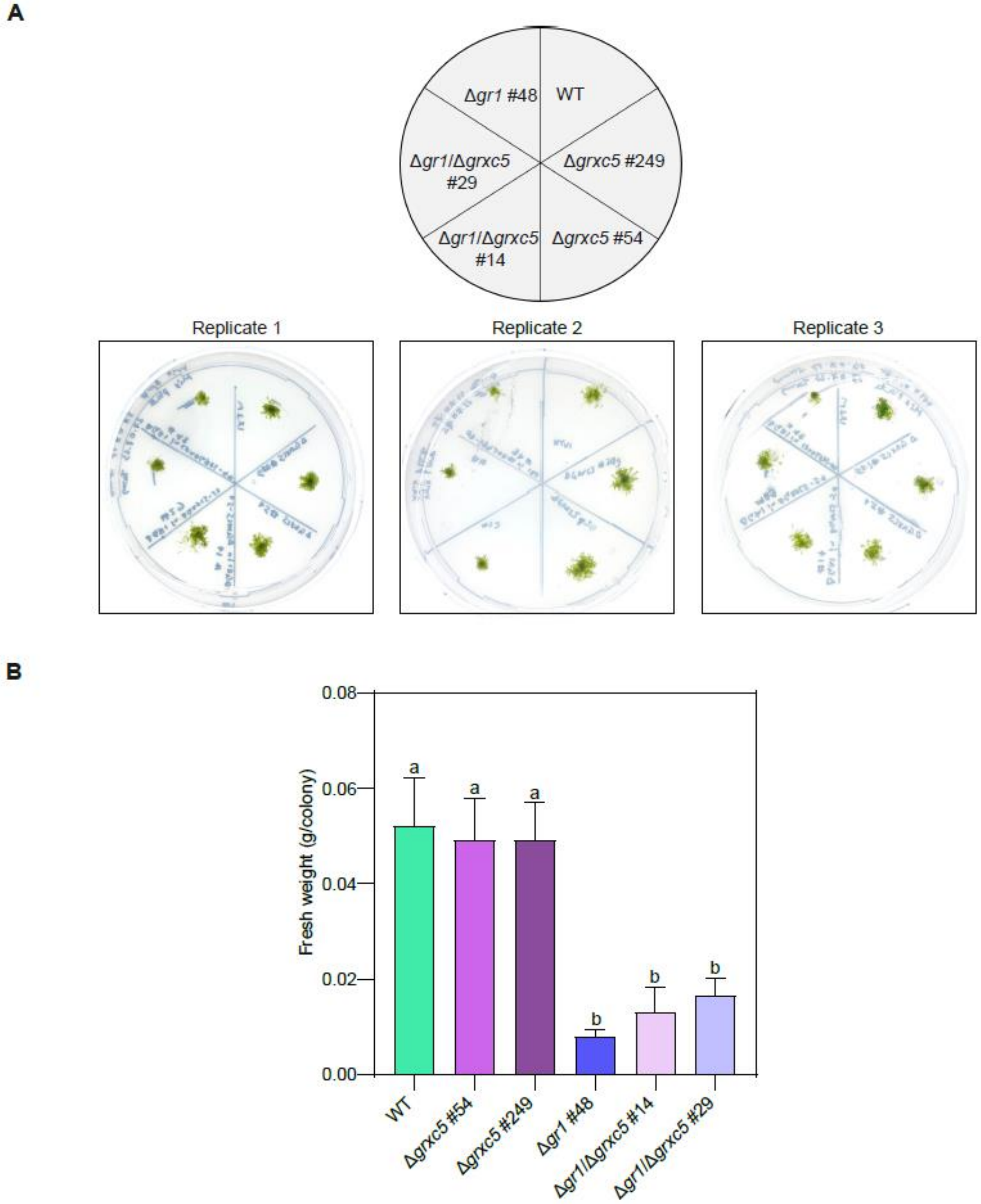


**Supplementary Figure S2: Fresh weight analysis for Δ*grxc5*, Δ*gr1* and Δ*grxc5*/Δ*gr1***
**(A)** *P. patens* grown on KNOP-ME pH 5.8 agar plates in 100 µmol photons m$^{-2}$s$^{-1}$ (16 h light, 8 h dark) for four weeks. **(B)** Fresh weight (g) of *P. patens* colony after four weeks (n=3; mean +SD). One-way ANOVA and Tukey's multiple comparison was performed to assess significant differences (p<0.0001) in fresh weight (n=3). Different lowercase letters indicate significant difference.

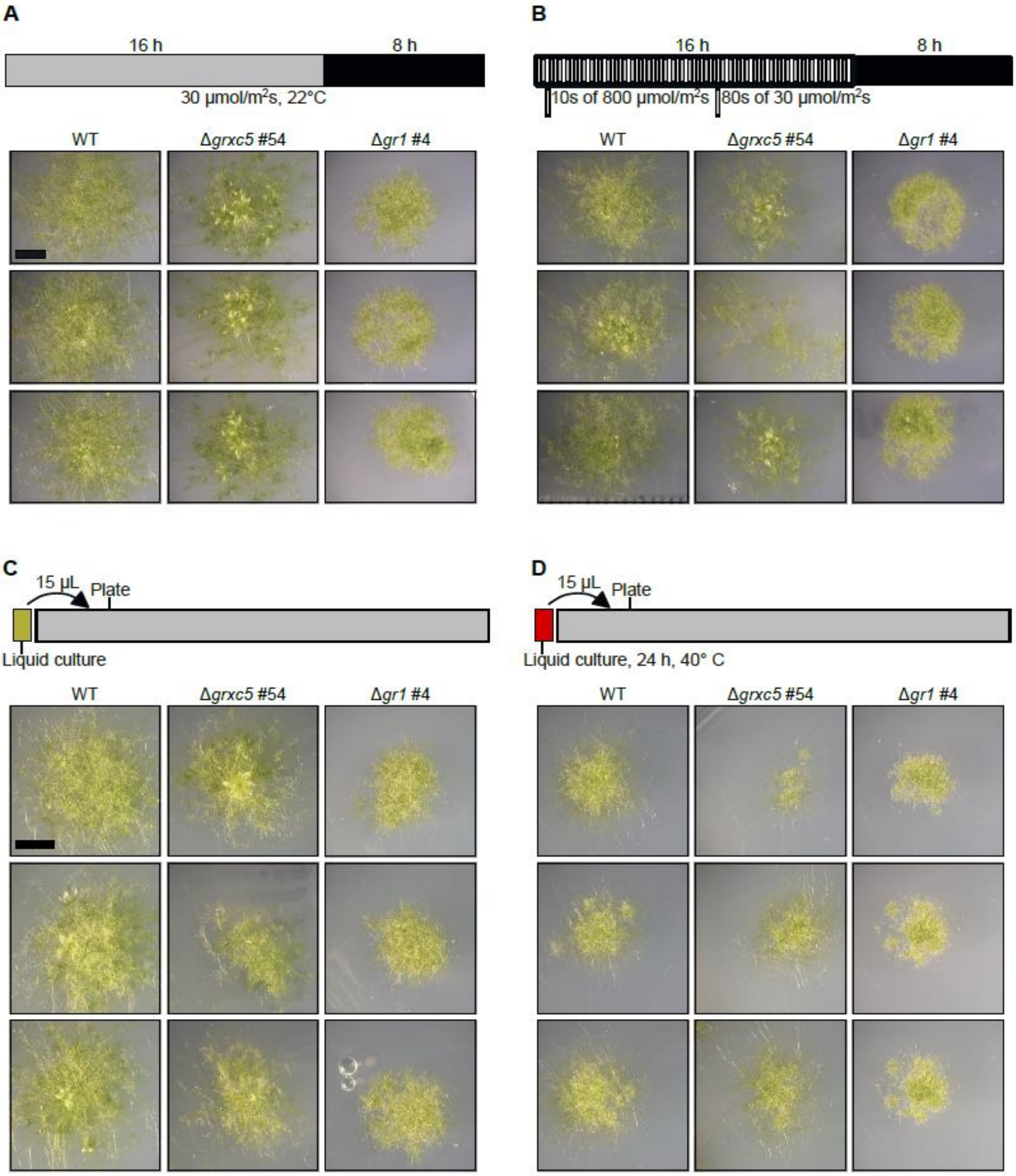


**Supplementary Figure S3: Stress phenotyping of *P.patens* protonema culture under fluctuating light and heat stress**

**(A, B)** Spotting assay using 15-20 µl of protonema culture on KNOP-ME plates after incubating under two different light conditions. In **(A)**, protonema was incubated under a constant light intensity of 30 µmol photons $m^{-2}s^{-1}$ for 14 d. In **(B)**, protonema was exposed to a fluctuating light regime with 10 s of 800 µmol photons $m^{-2}s^{-1}$ followed by 80 s of 30 µmol photons $m^{-2}s^{-1}$ (Vaseghi *et al.*, 2018), under long day conditions (16 h/8 h). Scale bar = 4mm. **(C, D)** Spotting assay using 15-20 µl of protonema culture on KNOP-ME plates. In **(C)**, protonema was grown without any treatment, while in **(D)**, protonema was subjected to 24 h of incubation at 40°C in the dark. Both samples were cultivated under standard conditions of 60 µmol photons $m^{-2}s^{-1}$, 22°C, with a 16-hour light/8-hour dark cycle. Images were taken after 20 d of growth. Scale bar = 4 mm.

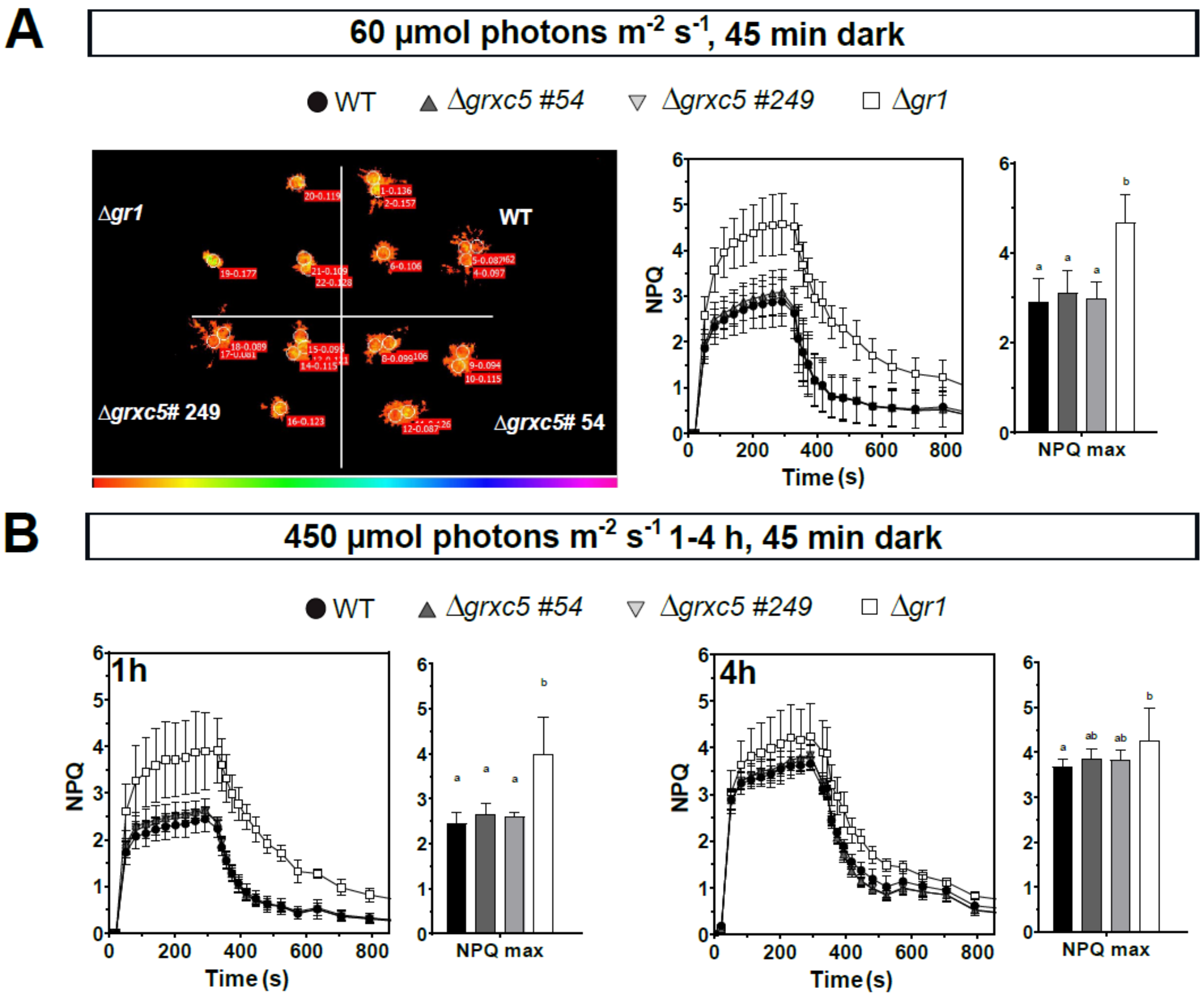


**Supplementary Figure S4: Non-photochemical quenching (NPQ) measurements of 4-week-old gametophores under low and high light regimes**

4-week-old gametophore colonies measured with an imaging PAM either grown in control light at 60 µmol photons $m^{-2}s^{-1}$ **(A)** or treated for 1 h or 4 h with light intensities of 450 µmol photons $m^{-2}s^{-1}$ prior to experiment start **(B)**. Plants were dark-adapted for 45 min before each measurement. Photosynthetically active radiance (PAR) was kept at 531 nm. Three agar plates containing three moss colonies of each plant line (n=9) were measured for control conditions, two agar plates for 1 h of 450 µmol photons $m^{-2}s^{-1}$ treatment (n=6) and one plate for 4 h 450 µmol photons $m^{-2}s^{-1}$ treatment (n=3) (each shown as mean + SD). Example overview of $F_0$ (dark-adapted) emitted by the moss colonies shown in A (left panel). A one-way ANOVA with Tukey's multiple comparison test was conducted on $NPQ_{max}$ to test for significant differences between the genotypes ($p<0.0055$). Small letter code: samples with different letters differ significantly.

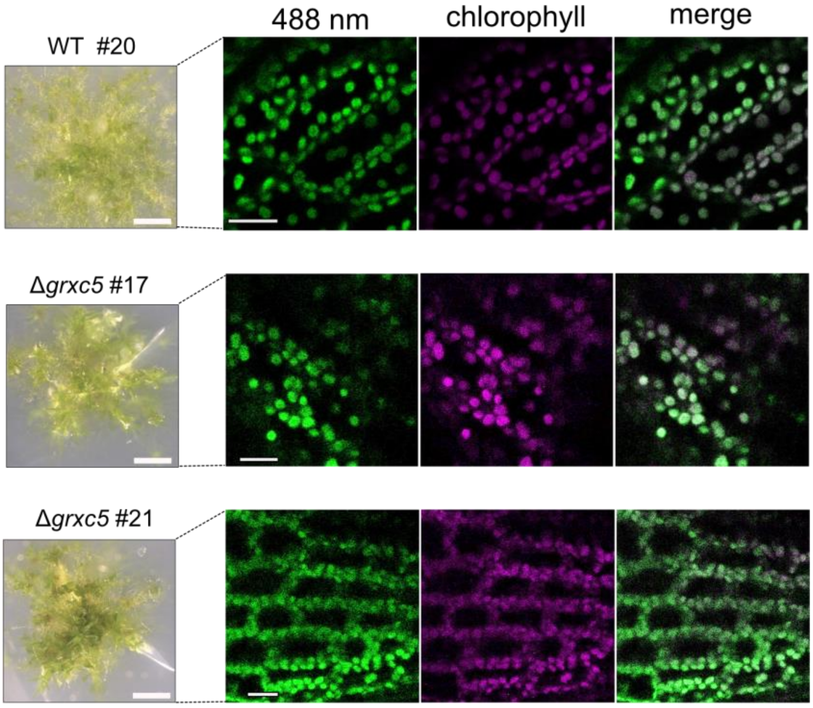


#### Supplementary Figure S5: Targeting of roGFP2 to plastids

**(Left)** Growth of 4-week-old *P. patens* gametophores on KNOP-ME under standard growth conditions. Scale bar = 0.2 mm. (**Right**) Example confocal images of *P. patens* expressing *TKTP-roGFP2*. roGFP2 was excited at 488 nm with emission recorded at 508-535 nm. Chlorophyll fluorescence was detected at 680-735 nm after excitation with 488 nm. Merge shows overlap of roGFP2 (488 nm) and chlorophyll signals. Scale bars = 20 µm.

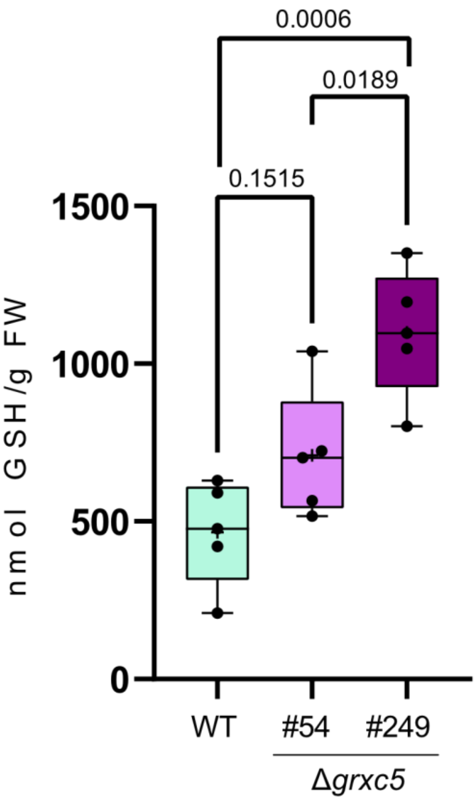


**Supplementary Figure S6: Quantification of total GSH in GRXC5 mutant lines by HPLC**
High-performance liquid chromatography (HPLC) measurements of total glutathione content (GSH + GSSG, n =5) of *P. patens* protonema grown in liquid medium was performed as described in Müller-Schüssele *et al.* (2020). Cultures were dispersed and transferred to fresh medium three days before harvest. Significant differences between biological replicates (one-way ANOVA, Tukey's multiple comparison) and respective p-values are depicted above the brackets. Whiskers show min and max values with the boxes display the 25-75 percentiles. The median is depicted as line.

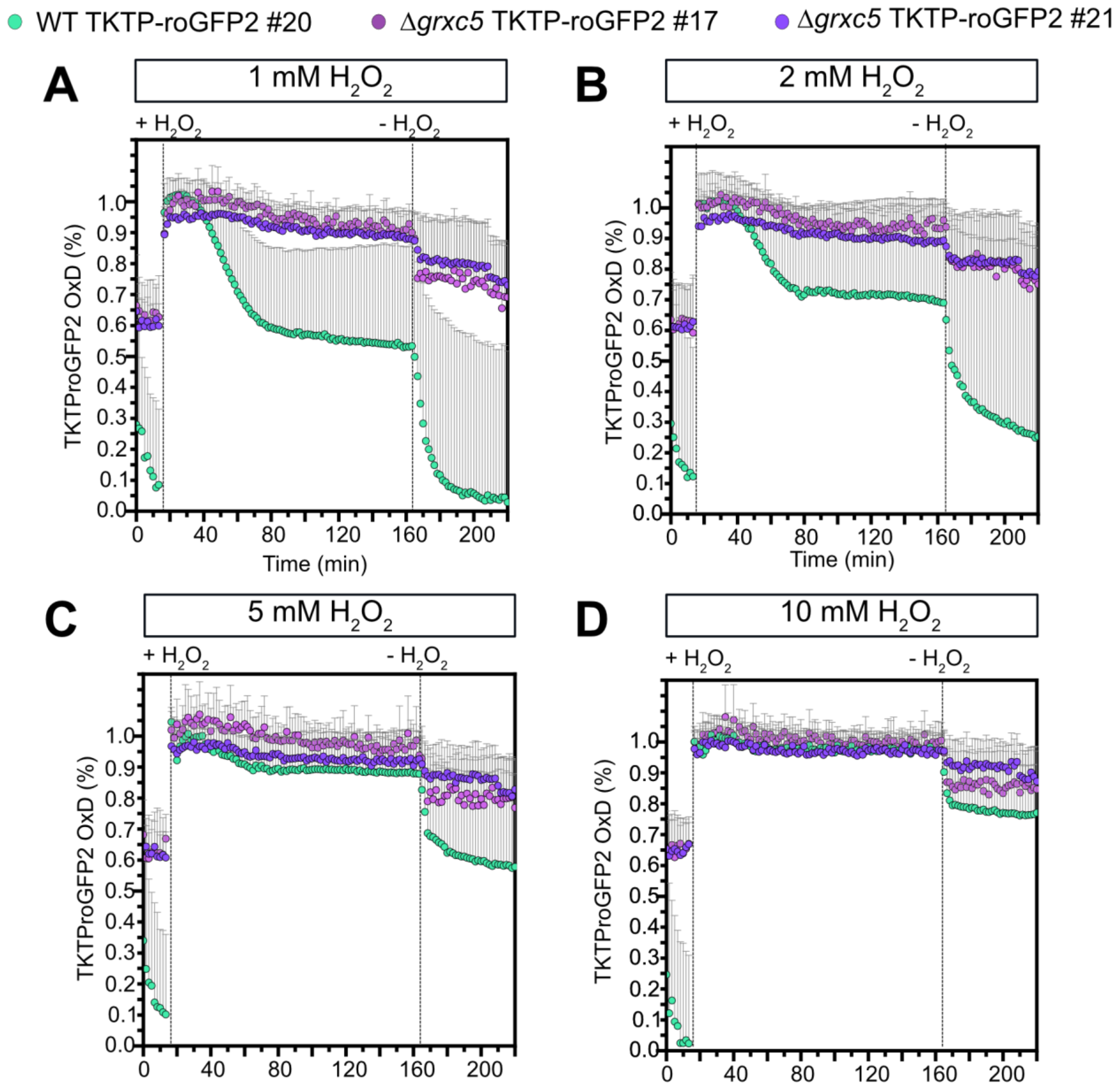


**Figure S7: roGFP2 oxidation state and recovery in response to externally added $H_2O_2$**
Protonema culture of *P. patens* expressing *TKTP-roGFP2* was imaged for 16 min before addition of 1 mM (**A**), 2 mM (**B**), 5 mM (**C**) 10 mM (**D**) $H_2O_2$. Addition or removal of $H_2O_2$ are indicated by dotted lines. For fully oxidized and fully reduced controls (calibration) 10 mM $H_2O_2$ or 10 mM DTT were preincubated for 30 min with the protonema culture and used for calculation of the degree of oxidation of roGFP2. After 165 min the buffer containing peroxide was exchanged to imaging buffer to monitor *in vivo* recovery of reduced roGFP2. Three biological replicates with three technical replicates each were conducted for each line and treatment (n=9). Shown are the mean + SD values.

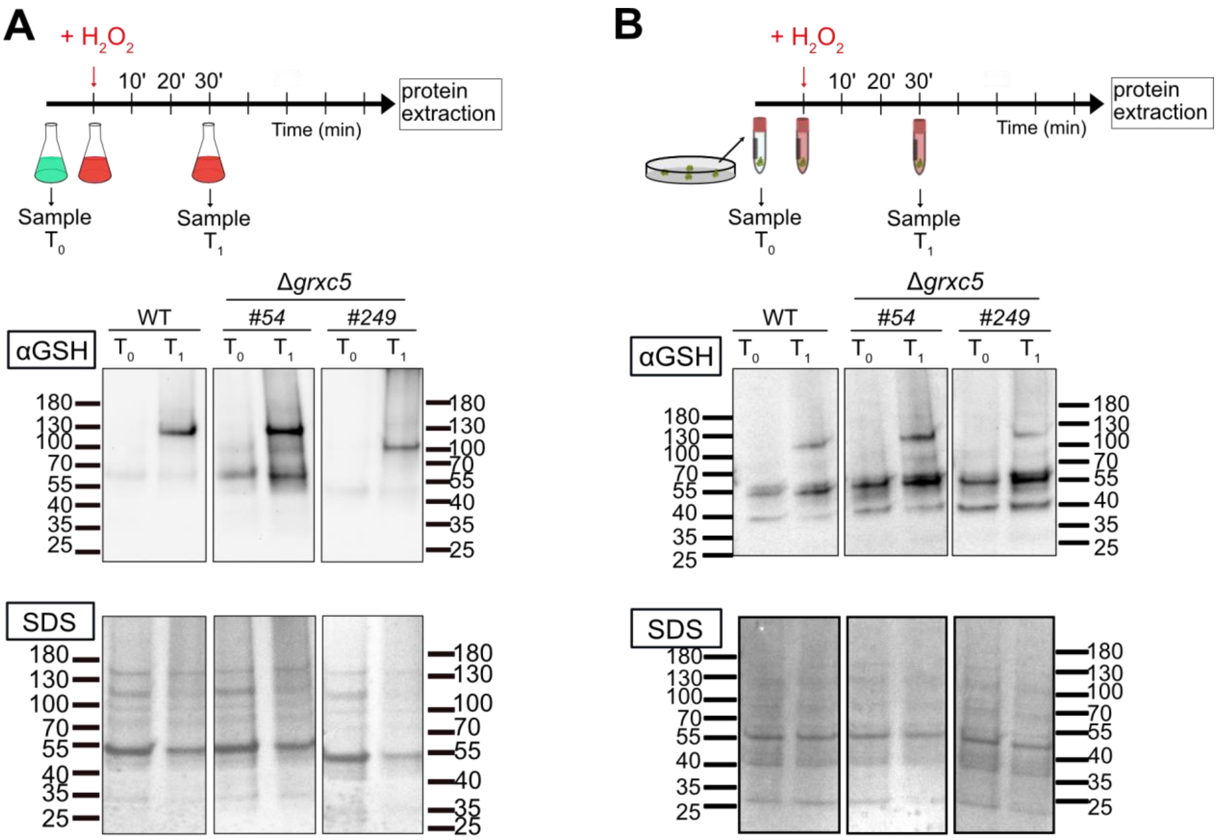


**Supplementary Figure S8: Increase of protein-bound glutathione after $H_2O_2$ treatment**
Schematic overview of the experimental set-up and immunodetection of protein-bound glutathione using proteins extracted (in the presence of 20 mM NEM) from *P. patens* protonema tissue **(A)** or gametophore tissue **(B)** treated with 10 mM $H_2O_2$ and harvested after 30 min ($T_0$ (before treatment), $T_1$ (30 min $H_2O_2$ incubation);10 µg protein extract was loaded (4-20% non-reducing SDS-PAGE).

**Supplemental Table 1: Primer sequences**

| Primer | Sequence | Purpose |
|---|---|---|
| **Triple template PCR** | | |
| PpGrxC5ko_5PHR P1 | ATCACAGGAAGCTATGGAAGGCA | Δ*grxc5* construct |
| PpGrxC5ko_5PHR P2 | TTGACAGGATCCGATAATCCCCACTTAGCA CCAGG | Δ*grxc5* construct |
| PpGrxC5ko_3PHR P3 | ATCGGGCCTCCTGTCATGCCATCACATACG GAACT | Δ*grxc5* construct |
| PpGrxC5ko_3PHR P4 | ATCTTCAGCTCCTCAGTTCCTCG | Δ*grxc5* construct |
| GrxC5ko_npt_F | TGCTAAGTGGGGATTATCGGATCCTGTCAA ACACTG | *nptII* resistance Δ*grxc5* construct |
| GrxC5ko_npt_R | CGTATGTGATGGCATGACAGGAGGCCCGA TCTAGTA | *nptII* resistance Δ*grxc5* construct |
| GrxC5_5P_F | AAGTAGGGAAAAGAGAGCACG | 5’ integration |
| H3b_R | CCAAACGTAAAACGGCTTGT | 5’ integration |
| NosT_F | GCGCGGTGTCATCTATGTTA | 3’ integration |
| GrxC5_3P_R | TGTCGTGTGTTCGGACTTCT | 3’ integration |
| PpGrxC5_RT_F | TTAATCGGCAGGTGTGTGGA | cDNA *GRXC5* |
| PpGrxC5_RT_R | AAAAGCTTCTTCACGCGCAT | cDNA *GRXC5* |
| PpEF1a_RT_F | CGACGCCCCTGGACATC | cDNA control |
| PpEF1a_RT_R2 | CATGTTGTCACCCTCGAACC | cDNA control |
| PpGR1_RT_F | CTATCGGGGCTGGTAGTGG | cDNA *GR1* |
| PpGR1_RT_R | GGCTGCTCTCAAAATCGTGT | cDNA *GR1* |
| PpGR1_5P_F | GAAGCACAACAAAGAGAGGCA | 5’ integration |
| PpGR1_3P_R | GGCTCATTTCCGAAAGCAGT | 3’ integration |
| TKTP_F | GGGGACAAGTTTGTACAAAAAAGCAGGCT ATGGCGTCTTCTTCTTCTCT | Stroma-targeted roGFP2 |
| TKTP_roGFP2_R | CCTCGCCCTTGCTCACCAGCGCAGTCTCA GTT | Stroma-targeted roGFP2 |

| | | |
|---|---|---|
| TKTP-roGFP2_F | ACTGAGACTGCGCTGGTGAGCAAGGGCGA GGAG | Stroma-targeted roGFP2 |
| roGFP2-attB2_R | GGGGACCACTTTGTACAAGAAAGCTGGGT CTTACTTGTACAGCTCGTCCATG | Stroma-targeted roGFP2 |

#### Supplementary Material and Methods

**Measurement of photosynthetic parameters (light induction and relaxation curves)**

For NPQ-measurements, three moss colonies of each, WT, Δ*grxc5* #54, Δ*grxc5* #249 and Δ*gr1* were grown on the same agar plate at 60 µmol photons $m^{-2}$ $s^{-1}$ (16 h light/ 8 h dark) for 4 weeks. Plates were either 45 min dark incubated and measured within an IMAGING-PAM (Maxi, Heinz Walz GmbH, Germany) or treated for 1 h and 4 h with high light (450 µmol photons $m^{-2}$ $s^{-1}$) and subsequently dark incubated for 45 min before measurement.

**Glutathione measurement by HPLC**

Total glutathione was determined as in Müller-Schüssele *et al.* (2020). Approximately 30 mg protonema tissue were extracted with 10-fold volume of 0.1 M HCl and centrifuged for 10 minutes at 4°C. Twenty-five µl of supernatant were neutralised with 25 µl 0.1 M NaOH and the thiols were reduced with 1 µl 0.1 M dithiothreitol for 15 min at 37°C in darkness. Ten µl 1 M Tris/HCl pH 8.0 and 35 µl water were added and the thiols were derivatised by 5 µl of 0.1 M monobromobimane (Thiolyte® MB, Calbiochem) in darkness for 15 min at 37°C. After adding 100 µl 9 % acetic acid and centrifugation at 4°C for 15 minutes the samples were separated on high-performance liquid chromatography (HPLC; Spherisorb™ ODS2, 250 x 4.6 mm, 5 µm, Waters, Eschborn, Germany) using a linear gradient from 4 to 20% of buffer A (90% methanol, 0.25% acetic acid, pH 3.9) in buffer B (10% methanol, 0.25% acetic acid, pH 3.9). The conjugates were detected fluorometrically with excitation at 390 nm and emission at 480 nm.
